# Typical development of the human fetal subplate: regional heterogeneity, growth, and asymmetry assessed by in vivo T2-weighted MRI

**DOI:** 10.1101/2025.10.06.680743

**Authors:** Andrea Gondová, Jennings Zhang, Sungmin You, Seungyoon Jeong, Milton O. Candela-Leal, Caitlin K Rollins, P. Ellen Grant, Hyuk Jin Yun, Kiho Im

## Abstract

The subplate (SP) is a transient fetal brain compartment supporting neuronal migration, axonal ingrowth, and early cortical activity, yet the dynamics of its regional development remain poorly understood in vivo. Using T2-weighted fetal MRI of 68 typically developing fetuses (22–32 weeks gestational age, GA), we developed a semi-automated pipeline to quantify regional SP morphology (thickness, surface area, and volume). SP characteristics scaled strongly with GA and residual brain volume and showed marked regional differences. After correcting for geometric confounds, regional variation of SP thickness persisted, with highest values in parietal and perisylvian regions, suggesting that SP thickness may serve as a sensitive marker of intrinsic developmental differences. Between late 2nd and early 3rd trimester, mean SP thickness increased by 39.2% with large variation across regions (±11.0 SD), whereas surface area growth was more uniform (64.3% ±0.7 SD). Continuous growth trajectories clustered into distinct spatiotemporal profiles: early-developing regions (e.g., pericentral and medial occipital cortices) contrasted with later-developing regions (prefrontal, temporal, and parietal cortices). These patterns partially recapitulate primary-to-association, medial-to-lateral, and posterior-to-anterior maturational hierarchies, pointing to organized developmental program. SP development also showed region-specific hemispheric asymmetries, including leftward thickness and volume asymmetry in superior temporal and precentral gyri. Some asymmetries amplified, others attenuated or reversed with age, suggesting both transient states and potential precursors of postnatal lateralization. Together, these findings provide a framework for regional SP quantification and position SP morphology, particularly thickness, as a promising early biomarker that might link fetal SP changes to subsequent cortical development and neurodevelopmental outcomes.

**Significance Statement:** The subplate (SP) is a transient fetal brain compartment that provides an early foundation for downstream cerebral development. Using fetal MRI, we show that SP morphology is regionally heterogeneous and that its growth follows distinct regional trajectories. Principal spatiotemporal patterns differentiate early-from later-developing regions and partially recapitulate sensory-to-association hierarchies of cortical maturation. SP also exhibits dynamic hemispheric asymmetries as early as the second trimester, with some amplifying as potential precursors of postnatal lateralization, and others reversing with age, emphasizing the importance of evaluating developmental trajectories rather than single time points. This framework positions SP morphology as an accessible early marker of fetal brain organization with potential relevance for later neurodevelopment.

## Introduction

Fetal brain development unfolds as a cascade of rapid and interrelated events that lay the foundation for mature brain organization and function^1,2^. The importance of this period for optimal neurodevelopment is increasingly recognized, based on accumulating evidence of lifelong functional consequences associated with adverse *in utero* environments or early-life events such as prematurity^2,3^ and *in utero* origins suggested for various neurodevelopmental disorders^4,5^.

A characteristic feature of the fetal brain from mid-gestation onward is the presence of transient compartments, most notably the subplate (SP)^6^. Foundational insights into SP development and function derive from animal models^7–11^ and *postmortem* human studies using histology and *ex vivo* magnetic resonance imaging (MRI)^6,12–15^. These investigations reveal SP’s complex architecture, including its sublaminar organization and the morphological and molecularly diverse populations of SP neurons, radial and tangential migratory streams, glial precursors, signaling molecules, and an abundant extracellular matrix (ECM)^16,17^. Crucially, the SP serves as a transient convergence zone for multiple afferent systems, including thalamocortical, basal forebrain, and early corticocortical projections^18–21^. The initial synaptic connectivity established within the SP contributes to early functional network activity^22,23^. At its peak, the SP can be up to four times thicker than the overlying cortical plate (CP), reflecting its high developmental load. As afferent progressively relocate to the CP, the SP gradually dissolves during the last third of gestation, marking a key transition toward more mature cerebral organization^24^. Crucially, the SP is not merely a transient compartment but actively participates in key corticogenic processes, shaping the formation of permanent neural circuits and cortical columnar organization by influencing CP differentiation and synaptogenesis. Through these roles, the SP likely contributes to cortical arealization and has been implicated in the mechanics of cortical folding^25^ (for review^16^). Thus, detailed *in vivo* characterization of SP morphology might offer a unique window into the early spatial patterning of the developing human brain, with important implications for understanding cortical developmental trajectories.

Importantly, atypical features of the SP have been hypothesized in several neurodevelopmental disorders, including cerebral palsy, autism spectrum disorder (ASD), attention deficit hyperactivity disorder (ADHD), and schizophrenia^26,27^. These underscore the potential of SP characteristics to serve as early biomarkers of later neurodevelopmental dysfunction. However, accurate interpretation of such deviations requires a detailed understanding of typical SP development, which itself is highly dynamic and marked by substantial spatial and temporal heterogeneity in its emergence, expansion, and dissolution^11^. This pronounced morphological variability is not only biologically significant but also presents unique computational challenges for SP quantification, which may partly explain why systematic *in vivo* assessment of the SP has been limited.

Increasingly, recent advances in *in utero* MRI, driven particularly by advances in rapid, motion-tolerant acquisition methods (e.g., HASTE^28^, echo-planar FLAIR^29^) and dedicated fetal brain processing pipelines^30–33^, are beginning to address these challenges, enabling *in vivo* study of the SP^34,35^. These technological advances have the potential to support anatomically precise cross-sectional as well as longitudinal investigations of fetal brain tissue compartments. The increasing availability of larger fetal datasets permits capturing developmental changes with higher temporal resolution. For example, several studies have quantified SP development using global metrics such as total volume^36,37^, T2 signal intensity^29^, and microstructural properties^38,39^. On a regional level, expansion of SP thickness has been described particularly in sensorimotor and occipital regions early in gestation^40^. Subsequent studies have extended these findings into later gestation, mapping regional SP volumes and linking them with sulcal emergence and gene expression gradients^41–43^.

However, gaps remain in this literature. Most prior *in vivo* studies have either focused on global SP metrics or included the subjacent intermediate zone (IZ), which limits regional and anatomical specificity. Although some studies have noted regional heterogeneity, few have assessed SP development using multiple morphological dimensions (e.g., thickness, surface area, volume), even though each dimension may exhibit distinct developmental signatures while reflecting interrelated biological processes. Moreover, SP assessment has been constrained by limited subject availability and also the time-consuming nature of manual segmentation or manual correction of registration-based SP labels. To address these limitations, we applied an in-house automated fetal brain processing pipeline, which enables robust segmentation of the SP and extraction of regional surface-based and volumetric features across the cerebrum^32,33,44^. This approach allows detailed in vivo characterization of SP development.

We focused on the 22–32 weeks gestational age (wGA) period. During this window, the SP appears as a distinct hyperintense band between the CP and IZ on T2-weighted MRI, primarily due to its ECM content^34^. Although the SP is histologically evident from 15–17 wGA and functionally significant by approximately 20 wGA^45,46^, earlier imaging is limited by small brain size and motion artifacts. By the upper limit of this window (~32 wGA), contrast between the SP and IZ progressively decreases due to ECM dissolution and afferent relocation to the CP, rendering the SP indistinguishable from the IZ^29,47^. Thus, the 22–32 wGA period offers optimal contrast for SP segmentation. Incidentally, this window also overlaps with peak SP prominence (before its progressive dissolution)^48^, coincides with intense proliferative and migratory events and axonal growth^48^, and aligns with the lower limit of viability in preterm birth^49^, making it both biologically and clinically relevant.

In this study, we characterized regional heterogeneity in SP thickness between 22–32 wGA to test whether inherent differences exist across the cerebrum. We then assessed regional SP development during this period by evaluating relative changes in regional SP thickness, surface area, and volume between the late 2^nd^ (22–27 wGA) and early 3^rd^ trimester (>27–32 wGA) periods, to identify regions of peak developmental activity. We also clustered regional developmental trajectories to identify common growth profiles. Finally, given the important hemispheric lateralization of adult brain organization^50–52^ and evidence for prenatal origins of this lateralization^39,53–58^ we examined interhemispheric asymmetries across regional SP characteristics to explore whether lateralized features were already apparent during this period and how they evolved with gestational age.

## Results

### Gestational age, brain volume, and regional variation modulate SP characteristics

Using a near-automated processing pipeline, we quantified SP thickness, surface area, and volume, enabling detailed assessment of developmental dynamics. First, we examined the global effects of GA, residual brain volume (supratentorial brain volume adjusted for GA), and sex on SP morphology. ANCOVA modelling revealed strong positive association with GA for whole-brain SP thickness (F=393.553, η^2^=0.774), surface area (F=6167.471, η^2^=0.950), and volume (F=2823.253, η^2^=0.927) (all p < 0.001; Supp. Table 2 & Supp. Fig. 5). Residual brain volume was independently associated with SP thickness (F=52.507, η^2^=0.103), surface area (F=248.569, η^2^=0.038), and volume (F=248.569, η^2^=0.052) (all p < 0.001), confirming scaling of SP morphology with overall brain size. Sex effects were minimal, likely because residual brain volume accounted for most of the sex-related variance, with only SP volume showing a significant but negligible sex effect (F=5.594, p=0.021, η^2^=0.002), and interactions with GA were also negligible (F=13.120, p=0.001, η^2^=0.002). Hence, sex was not considered in further analyses.

We next examined thickness variation across SP regions and hemispheres using repeated-measures ANCOVA. Cortical folding depth (‘SP depth’) is geometrically related to SP thickness, confirmed by a significant within-subject effect (F=190.129, p<0.001, η^2^=0.016) and a modest interaction with region (F=2.321, p=0.002, η^2^=0.003) (full statistical results are provided in Supp. Table 4). Substantial regional differences in SP thickness persisted after correcting SP thickness for regional SP depth to account for these geometric influences of outer SP undulations on thickness (F=111.956, p<0.001, η^2^=0.271; Figure 2a, Supp. Table 5). Small but significant interactions were also observed for region × GA (F=9.052, p<0.001, η^2^=0.022), region × hemisphere (F=26.278, p<0.001, η^2^=0.025), and their three-way interaction (F=4.215, p<0.001, η^2^=0.004), suggesting that growth rates of SP thickness vary regionally and show hemispheric asymmetry. These results indicate that SP thickness heterogeneity cannot be explained solely by global brain growth or differences in cortical folding, and may reflect regional differences in axonal ingrowth, cellular composition, or extracellular matrix volume.

Similar modelling of SP surface area and volume reveal small but significant interactions for region × GA (surface area: F=5.550, p<0.001; volume: F=23.520, p<0.001) and region × hemisphere (surface area: F=26.780, p<0.001; volume: F=29.731, p<0.001), along with their three-way interactions (surface area: F=3.180, p<0.001; volume: F=6.808, p<0.001), though effect sizes were small (η^2^<0.009).These interactions indicate subtle region- and hemisphere-specific growth patterns, but interpretation of absolute biological differences is limited, as the size of parcellated regions depend on the predetermined parcellation scheme. Overall, these analyses demonstrate that SP growth is closely coupled to overall brain development and size, with considerable regional and hemispheric variability, and suggest that thickness may provide a sensitive measure of region-specific heterogeneity, motivating subsequent analyses of regional growth patterns and hemispheric asymmetries.

### Analysis of regional SP growth identifies distinct spatiotemporal developmental patterns across 22-32wGA

We next characterized developmental dynamics of SP growth as a relative regional growth between late 2^nd^ (22-27wGA) and early 3^rd^ (27-32wGA) trimester, after adjusting regional values for same confounders as before within groups. SP thickness, surface area, and volume increased across all regions, but the extent of this growth varied by metric and region Figure 2b,c, Supp. Table 7). SP volume showed a mean growth of 81.3% and a large standard deviation (SD) of 16.6. The inferior temporal, fusiform, and middle frontal gyri showed the largest volume increases (up to +98%). In contrast, surface area growth was more uniform with a low SD (mean growth 64.3% ± 0.7), peaking in the lingual gyrus (+65%). Thickness changes were also heterogeneous (mean growth 39.2% ± 11.0 SD), with the greatest increases in the paracentral lobule, right superior parietal cortex, and left precentral and lingual gyri (+49%). These patterns suggest that SP growth is regionally orchestrated, with certain regions undergoing more rapid expansion during mid-gestation, potentially supporting early cortical circuit establishment.

To elucidate dominant spatiotemporal patterns, Ward’s hierarchical clustering was applied to continuous regional growth profiles. Four clusters were identified for SP thickness (Silhouette=0.513), and three clusters each for surface area (Silhouette=0.406) and volume (Silhouette=0.361) based on Silhouette scores and dendrogram inspection (Figure 3, Supp. Fig. 8, Supp. Table 8). Cluster-averaged profiles revealed differences in peak growth rate, timing, and cumulative growth (Table 1a).

**Table 1.** **a**, Summary of developmental trajectories for subplate (SP) clusters. For each cluster, the table reports peak relative growth rate (%/week), peak timing (wGA), cumulative growth (AUC, computed as the area under the relative growth rate curve), and growth density (standard deviation of the smoothed growth curve). **b**, Matrices comparing bilateral asymmetry across clusters. Each cell shows the normalized symmetry between left and right hemisphere regions within or across clusters (higher values indicate greater symmetry). Diagonal entries thus reflect the degree of symmetry for homotopic regions assigned to the same cluster.

| **a,** |  | *Peak rate (%/week)* | *Peak time (week)* | *AUC (%)* | *Growth density (%/week)* |
| --- | --- | --- | --- | --- | --- |
| **SP thickness** |  |  |  |  |  |
|  | Cluster T1 | 249.48 | 22.05 | 119.75 | 4.83 |
|  | Cluster T2 | 160.61 | 22.05 | 87.03 | 3.03 |
|  | Cluster T3 | 110.01 | 31.35 | 73.44 | 3.73 |
|  | Cluster T4 | 88.78 | 22.05 | 50.39 | 1.74 |
| **SP surface area** |  |  |  |  |  |
|  | Cluster SA1 | 139.35 | 30.25 | 91.99 | 5.37 |
|  | Cluster SA2 | 133.30 | 29.75 | 98.07 | 4.16 |
|  | Cluster SA3 | 129.13 | 29.65 | 98.88 | 3.61 |
| **SP volume** |  |  |  |  |  |
|  | Cluster V1 | 268.54 | 22.05 | 116.38 | 5.52 |
|  | Cluster V2 | 139.17 | 27.75 | 121.09 | 2.19 |
|  | Cluster V3 | 150.06 | 29.35 | 107.92 | 5.15 |

For instance, SP thickness clusters T1 and T2 peaked early: T1 (e.g., right superior frontal, bilateral paracentral lobule, lingual gyri) showed the highest peak rate (249.5%/week) and cumulative growth (AUC=119.8%), while T2 (areas around central sulcus, left temporal lobe) had more moderate growth density (i.e., temporal concentration of growth; 3.03%/week) and cumulative growth (AUC=87.0%). T3 (mainly parietal and parts of temporal cortex) peaked the latest (~31.35 wGA) with slower growth (110%/week). T4 (lateral occipital, and parts of right temporal lobe, precuneus) peaked early but showed the lowest growth density (1.74%/week) and cumulative growth (AUC = 50.4%). Surface area clusters peaked more uniformly around 30 wGA. The primarily frontal cluster SA1 showed highest growth density (5.37%/week) and latest peak (30.25 wGA), while SA2 and SA3 showed similar cumulative growth (AUC~98%) with lower growth density (<4.16%/week). Volume clusters displayed more heterogeneity: V1 peaked early with the highest growth rate (268.5%/week) and growth density (5.52%/week); V2 (mainly parietal) peaked later (27.75w) with the highest cumulative growth (area under the relative growth curve [AUC] = 121.1%) and moderate growth density (2.19%/week); V3 (mainly frontal) peaked last (29.35w) with high growth density (5.15%/week) but lower cumulative growth (AUC = 107.9%).

To summarize the main patterns, early-peaking clusters (e.g., T1/T2, V1) exhibited rapid, temporally concentrated growth compared to later-peaking clusters (e.g., T3, SA1–3, V3) with delayed expansion. Some thickness regions (T4) exhibited flatter trajectories, suggesting sustained but low-intensity growth. These temporal and spatial patterns may reflect coordinated sequences of SP development (similar developmental sequences were previously suggested for maturation of white matter connectivity and cortical microstructure^59^) and could provide quantitative insights into the typical patterns and timing of early developmental events across cortical regions.

### SP characteristics show robust, region-specific hemispheric asymmetries that evolve between late 2^nd^ (22-27wGA) and early 3^rd^ (27-32wGA) trimester

Comparisons of homotopic regions for SP thickness, surface area, and volume with paired t-tests revealed significant left-right differences after adjusting for confounders (GA, residual brain volume, and additionally regional SP depth in case of SP thickness), with asymmetry indices (AIs) confirming consistent lateral%ization patterns (Figure 4a). Rightward asymmetries were observed in middle frontal, inferior frontal, and postcentral gyri across all SP characteristics (T=3.794–13.519, Cohen’s D=0.28–0.61, p<0.001– 0.01, FDR-corrected across regions within metric). Additional rightward asymmetries were seen for example in lingual gyrus and superior parietal cortex (thickness, volume), fusiform gyrus (thickness), and paracentral lobule (volume). The precuneus consistently showed leftward asymmetry across all metrics (T=–14.386 to –4.391, Cohen’s D=0.42–1.59, p<0.001). Leftward asymmetries were also present in cuneus and inferior parietal cortex (volume and thickness), precentral gyrus (volume), and supramarginal gyrus (surface area). Some regions, including lateral occipital cortex, superior frontal, and middle temporal gyri, exhibited metric-specific asymmetries, highlighting that thickness, surface area, and volume each capture distinct aspects of SP development.

Analysis of changes in AIs over time revealed that asymmetries can emerge, amplify, attenuate, or even reverse in a region-specific manner between late 2^nd^ and early 3^rd^ trimester (Figure 4b, Supp. Table 7). For example, the lingual gyrus showed a significant increase in rightward SP volume asymmetry (ΔAI = 5.08, Z = 2.35, p = 0.036), while the lateral occipital cortex, paracentral lobule, and postcentral gyrus developed new rightward asymmetry by mid-gestation. The precuneus maintained strong leftward asymmetry with a significant volume AI increase (ΔAI = 5.12, Z = 2.46, p = 0.035). Dynamic changes included attenuation of leftward volume asymmetry in the inferior parietal cortex due to rightward-biased growth (ΔAI = 6.07, Z = 2.40, p = 0.035), reversal of the superior frontal gyrus from leftward to rightward asymmetry (late 2^nd^: AI=–4,18, p=0.043; early 3^rd^: AI=3.13, p=0.003), and reversal of thickness asymmetry in the inferior temporal gyrus from rightward to leftward (late 2^nd^: AI=10.27, p=0.006; early 3^rd^: AI=−4.19, p=0.048).

Cluster spatial analysis indicated moderate bilateral symmetry across SP characteristics, strongest in volume clusters and weakest in thickness clusters, consistent with thickness’s greater variability in its spatial and temporal developmental profiles (Table 1b).

In summary, SP development shows early hemispheric asymmetries identifiable across late 2^nd^ and early 3^rd^ trimesters that are both dynamic and region-specific: some likely transient, others potentially providing a basis for postnatal lateralization. While regional comparisons reveal clear asymmetries, the underlying growth profiles are largely symmetric, indicating that lateralization might arise from subtle deviations within an otherwise coordinated bilateral program.

## Discussion

We present a framework for in vivo quantification of regional SP morphology using T2-weighted fetal MRI, enabling assessment of thickness, surface area, and volume between 22–32 wGA. Our findings demonstrate that SP morphology is both regionally heterogeneous and develops along region-specific trajectories. Relative growth analyses and hierarchical clustering reveal principal spatiotemporal patterns, differentiating early-from later-developing regions. SP also exhibits dynamic, region-specific hemispheric asymmetries emerging as early as the second trimester. Together, these findings highlight the complex interplay of intrinsic regional properties, growth timing, and lateralization, underscoring SP’s role as an early scaffold for downstream cortical development.

### Regional heterogeneity in SP thickness

SP thickness showed pronounced regional differences that persisted after controlling for GA, residual brain size, and SP depth. Parietal and perisylvian regions exhibited the greatest thickness, consistent with histological and imaging studies^21,34,40,63^. These differences likely reflect multiple factors, including differential afferent input, for example, early thalamocortical and dense associative connectivity linked to higher SP thickness in perisylvian and parietal areas^11,45,64^ or higher somatosensory thalamic^21^ compared to motor afferents that could explain greater SP thickness in postcentral vs precentral gyrus. Intrinsic regional factors, including previously described variations in neuron density, molecular phenotype, glial composition, and ECM also likely further module thickness directly or indirectly through further modulation of afferent targeting and synaptic remodeling^16,17,24,66^ and merit further study to elucidate the drivers of inherent differences in SP thickness across regions. Studies using diffusion imaging, particularly in conditions altering afferent input (e.g., thalamic injury, callosal agenesis) will be useful in this context.

Importantly, our findings complement prior work showing spatial patterning of other transient fetal compartments^1^. Combined with previous observations of consistent regional CP/SP volume proportions across gestation, and links between CP and SP thickness and regional gene expression^42,67^, the evidence is consistent with the protomap hyporthesis, suggesting that fundamental organization and regional identity may be determined early in development by intrinsic factors^70,71^. Interestingly, SP is not a passive relay but an active scaffold for early circuit formation. For example, SP neurons serve as pioneers scaffolding axonal pathways^68^, with specialized SP corridors guiding thalamic axons described in occipital regions^69,^ and SP further influences cortical plate differentiation and synaptogenesis^22,72,73^. Although we project cortical regions onto the underlying SP assuming a one-to-one correspondence, this relationship is likely more dynamic, shaped by tangential migration and regional developmental gradients^74^ rather than a fixed columnar organization, *in vivo* SP thickness may serve as a relatively accessible quantifiable marker of early arealization with potential to predict later cortical development that deserves further attention.

### Asynchornous regional growth between late 2^nd^ and early 3^rd^ trimester

Our results demonstrate asynchronous regional SP development, echoing prior histological and imaging work^75^. Between late 2^nd^ and early 3^rd^ trimesters, relative growth in SP thickness varied widely: paracentral lobule showed the greatest increases, potentially reflecting the changes in its motor-related part aligning with late second trimester motor area maturation^75^. Precentral gyrus thickening aligned with this in comparison to postcentral gyrus which showed more modest change, possibly due to earlier somatosensory afferent arrival. Medial occipital regions (cuneus, lingual gyrus) thickened more than lateral occipital cortex, consistent with earlier medial visual system maturation^67^. Association cortices such as inferior parietal and temporal areas, linked to human-specific evolutionary expansion^76,77^, also exhibited pronounced thickening, perhaps reflecting prolonged SP development in evolutionarily significant areas.

In contrast, surface area growth was more uniform, suggesting broadly isometric surface expansion during this gestational window. We speculate that global tangential expansion might be spatially constrained by adjacent regions, favoring greater radial thickening in areas with pronounced SP growth, particularly in association cortex, to mitigate developmental demands and spatial competition. Such localized thickening may in turn alter regional biomechanics, potentially influencing downstream folding patterns, especially relevant to secondary and tertiary sulci that emerge after 32wGA^78^ and to the expanded surface area of association cortex^76^. Prior work implicates subcortical stiffness and growth gradients in folding mechanics^79,80^, and imaging evidence suggests that SP changes precede sulcal formation^38,81^. Whether regional SP thickening modulates local stress and contributes to species-specific folding remains unresolved and will require integration of high-resolution fetal diffusion MRI, biomechanical modeling, and comparative developmental studies.

### Dominant growth profiles from clustering

Hierarchical clustering of continuous regional trajectories revealed dominant developmental patterns for thickness, surface area, and volume. Early thickening clusters (T1: paracentral lobule, lingual gyrus; T2: pericentral, cuneus, left temporal regions) partially align with early sensory systems^13,82^, while later clusters (T3: parietal and lateral temporal gyri) followed more delayed thickening potentially typical of association cortex^16^. These patterns partially recapitulate sensory-to-association hierarchies observed in white matter connectivity or cortical microstructure^59^, and may anticipate postnatal sensory-association gradients of functional organization^83–85^. SP volume clusters showed similar ‘gradient’ (early-growing V1: postcentral, middle temporal; followed by later, more steadily growing V2: parietal, lateral occipital, inferior temporal; and V3: frontal, medial occipital, superior temporal). Exceptions, such as the left middle temporal gyrus grouping with early-thickening regions, may reflect accelerated development of its posterior, more sensory-related subregions^86^, underscoring the limitations of our relatively coarse parcellation scheme which might mask finer-scale heterogeneity. Differences between medial vs. lateral occipital regions (T1/T2 vs T3), and paracentral vs. pre/postcentral gyri (T1 vs T2) further suggest spatially organized development, suggestive of lateral-to-medial (and potentially posterior-to-anterior) sequences akin to those in other developing tissues ^86^. Overall, our results support spatially organized developmental programs within SP that reflect known developmental patterns and could anticipate future functional organization while interacting with asynchronous, region-specific processes, highlighting SP development as a critical entry point for understanding cortical development.

### Asymmetries of SP morphology and their developmental dynamics

Hemispheric asymmetry is a known feature of mature brain organization^51,52^, with some prenatal origins suggested by prior imaging^53,55,56^. However, description of such asymmetries in transient fetal structures like the SP remains limited. Here, we report significant region-specific lateralization across SP characteristics. For example, the superior temporal gyrus showed leftward thickness and volume asymmetry, aligning with prenatal left temporal enlargement^53^, while rightward surface asymmetry in superior and middle temporal gyri paralleled earlier right sulcal development^55^. Precentral and postcentral gyri displayed opposing volume asymmetries: leftward in precentral (mirroring adult motor asymmetry^51^) and rightward in postcentral, the latter potentially novel observation. Frontal regions were mostly right-lateralized except the inferior frontal gyrus, partly diverging from previous fetal studies^56^. Such variation likely reflects the individual heterogeneity^87^, dynamic nature of SP development, and methodological limitations. For example, although we used a symmetric parcellation scheme, residual asymmetries may partly reflect parcel misalignment due to folding asymmetries, which could influence some region-specific estimates.

Additionally, asymmetry profiles evolved over gestation. Some regions (e.g., lingual gyrus, precuneus, pre/postcentral gyri) maintained or strengthened asymmetry, suggesting early subtle biases, including those in regional gene expression and molecular signatures, amplified over time^88^. In contrast, other asymmetries attenuated or reversed during the evaluated period (e.g., inferior parietal cortex showing leftward volume but decreasing over time; attenuation of surface asymmetry in inferior temporal gyrus and paracentral lobule; or rightward-to-leftward reversal of thickness asymmetry in inferior temporal gurus and the reverse for volume asymmetry in superior frontal gyrus). These findings indicate that while some fetal SP asymmetries resemble patterns observed later in life and could represent early precursors of postnatal lateralization, their dynamic nature highlights the complexity of fetal SP lateralization. This cautions against direct extrapolation to adult patterns, particularly given extensive sensory-dependent perinatal cerebral remodeling, and underscores the importance of both timing assessments and evaluating developmental trajectories over extended periods.

Based on developmental profiles, homotopic regions largely clustered together for volume and surface area growth, and to a lesser degree for thickness, indicating symmetrical timing and growth dynamics despite morphometric asymmetry. We propose that this suggests that SP asymmetries arise from localized regional differences^88^ superimposed on broadly symmetrical developmental programs. Given SP’s role in afferent integration and circuit scaffolding, early lateralization may bias later functional specialization, and its disruption could contribute to neurodevelopmental disorders in which altered asymmetries have been reported (e.g., autism, ADHD, dyslexia) ^51,89,90^. Determining which fetal asymmetries persist and which resolve will be essential for understanding the origins of these altered patterns, making longitudinal, multimodal studies of SP morphology particularly valuable.

### Conclusion

Our *in vivo* characterization of SP between 22–32 wGA reveals spatially heterogeneous, temporally dynamic, and asymmetrically patterned development. Regional trajectories of thickness, surface area, and volume, along with clustering analyses, identify spatially organized programs that partially recapitulate known developmental hierarchies. Early, evolving asymmetries further demonstrate fetal lateralization. These results establish a robust framework for in vivo SP quantification, offering an early entry point to study how fetal SP development influences downstream cortical maturation, connectivity, folding, and neurodevelopmental outcomes, providing a foundation for future biomarker studies.

## Materials and Methods

### Subjects

This study included fetal MRI collected at Boston Children’s Hospital from 2014 to 2024 under IRB approval (IRB-P00008836). Written informed consent was obtained from all parents. Fetuses with congenital brain malformations, incidental MRI findings, or serious maternal medical conditions were excluded. The final cohort included 68 typically developing singleton fetuses (34 males, 32 females, 2 unknown sex), scanned at mean 27.4wGA, range: 22.0-31.4wGA (Supp. Fig. 1). Only scans before 32 wGA were included because SP contrast becomes indistinguishable from the intermediate zone on T2-weighted MRI beyond this age^29^, as confirmed in our data by SP/IZ contrast approaching 0 towards 32wGA (Supp. Fig. 2). Future multimodal imaging incorporating complementary T1- and T2-weighted data^37^, unavailable in our dataset, may extend these observations into late gestation. Mean maternal age was 32.6 years, range: 21.8–40.0. For developmental analyses, fetuses were grouped into late 2^nd^ (22–27 wGA,N=29) and early 3^rd^ ((>27– 32wGA, N=39) trimester groups. A summary of cohort characteristics is provided in Supp. Table 1.

### Data acquisition and reconstruction

MRI was performed on 3T scanners: 42 scans on a Siemens MAGNETOM Skyra and 26 on a Siemens MAGNETOM Prisma (Siemens Healthineers, Germany). Imaging included repeated multi-planar T2-weighted half-Fourier single-shot turbo spin echo (HASTE) sequences optimized for fetal imaging. For each subject, HASTE stacks were acquired in axial, coronal, and sagittal orientations at least three times, totalling ~30 minutes of acquisition time. Parameters included TR = 1400–1600 ms, TE = 99–132 ms, 1 mm in-plane resolution, 2–3.5 mm slice thickness, and variable field of view adapted to fetal and maternal size^91^. Data was pre-processed with a dedicated pipeline that included brain extraction^44^, N4 bias field correction^92^, and slice-to-volume reconstruction with NeSVoR^32^, resulting in super-resolved motion-corrected 3D T2w volumes at 0.5 mm isotropic resolution.

### SP Segmentation and surface extraction

T2w volumes were registered to a 31w fetal brain template using rigid-body registration with scaling (7 dof) with FSL’s FLIRT^93,94^, as this template space is expected by our segmentation models. SP segmentation was performed with an ensemble of 2D attention-gated U-Net models trained on sagittal, coronal, and axial views, with aggregation and test-time augmentation to improve robustness. This approach adapted a previously validated CP segmentation method^44^ fine-tuned to include SP labels^60^. Segmentations were visually inspected and manually corrected by a single trained rater to address any segmentation errors based on underlying T2-weighted images. Although no formal reliability testing was performed, the SP’s simple morphology and the minimal manual edits required suggest the segmentations were sufficiently accurate for regional volume assessments.

Outer SP surfaces (CP/SP boundary) were extracted using the CIVET marching cubes algorithm^95^ using the ChRIS pl-fetal-surface-extract plugin^96^ and post-processed with Taubin smoothing^97^ to suppress overfitting to the coarse voxel boundaries. Inner surfaces (SP/IZ boundary) were generated by inward deformation along subject-specific radial distance maps optimized according to subject’s estimated gyrification index with empirically determined adaptive scheduling to accommodate diverse developmental morphologies^98^. This approach thus preserves mesh topology and vertex correspondence between outer and inner SP surfaces. Surfaces were resampled to standardized meshes of 81,920 triangles and 40,962 vertices. Surface quality was evaluated visually and quantitatively via smoothness error (mean curvature difference between each vertex and its neighbors) and boundary distance error (Euclidean distance of vertex to nearest boundary voxel) (Supp. Fig. 3).

### Regional parcellation

21 bilateral regions were delineated by adapting original 34-region Desikan-Killiany atlas^99^ to the fetal brain as proposed in^41^ with three exceptions: (1) separate superior, middle, and inferior temporal gyri; (2) subdivision of pericalcarine into cuneus and lingual gyrus; and (3) merging gyrus rectus with orbital frontal gyrus (Figure 1b, Supp. Table 2). Parcellations were manually defined on a 29w surface template^94,100^ and projected to individual SP surfaces via 2D spherical warping^101,102^. Surface labels were then mapped to volumetric space using a ribbon-constrained projection adapted from Connectome Workbench (v2.0.1), restricted to the SP label to yield volumetric SP regions. Regions with poor data quality were excluded: orbitofrontal cortex (due to frequent fronto-ventral blur on T2w), and peri-cingular pole regions: cingulate gyrus, insula, and parahippocampal gyrus (due to surface extraction artifacts caused by abrupt SP label termination leading to local over- or under-estimation of surface geometry to avoid propagating errors in measurements), resulting in 17 bilateral regions. Nevertheless, while our parcellation scheme was adapted for fetal anatomy^67^, it is ultimately based on adult cortical landmarks^99^, which may not perfectly correspond to fetal topography, especially before sulcation. This mismatch, together with potential registration inaccuracies, particularly in regions lacking clear anatomical landmarks, may reduce spatial precision and affect regional specificity.

**Figure 1.**
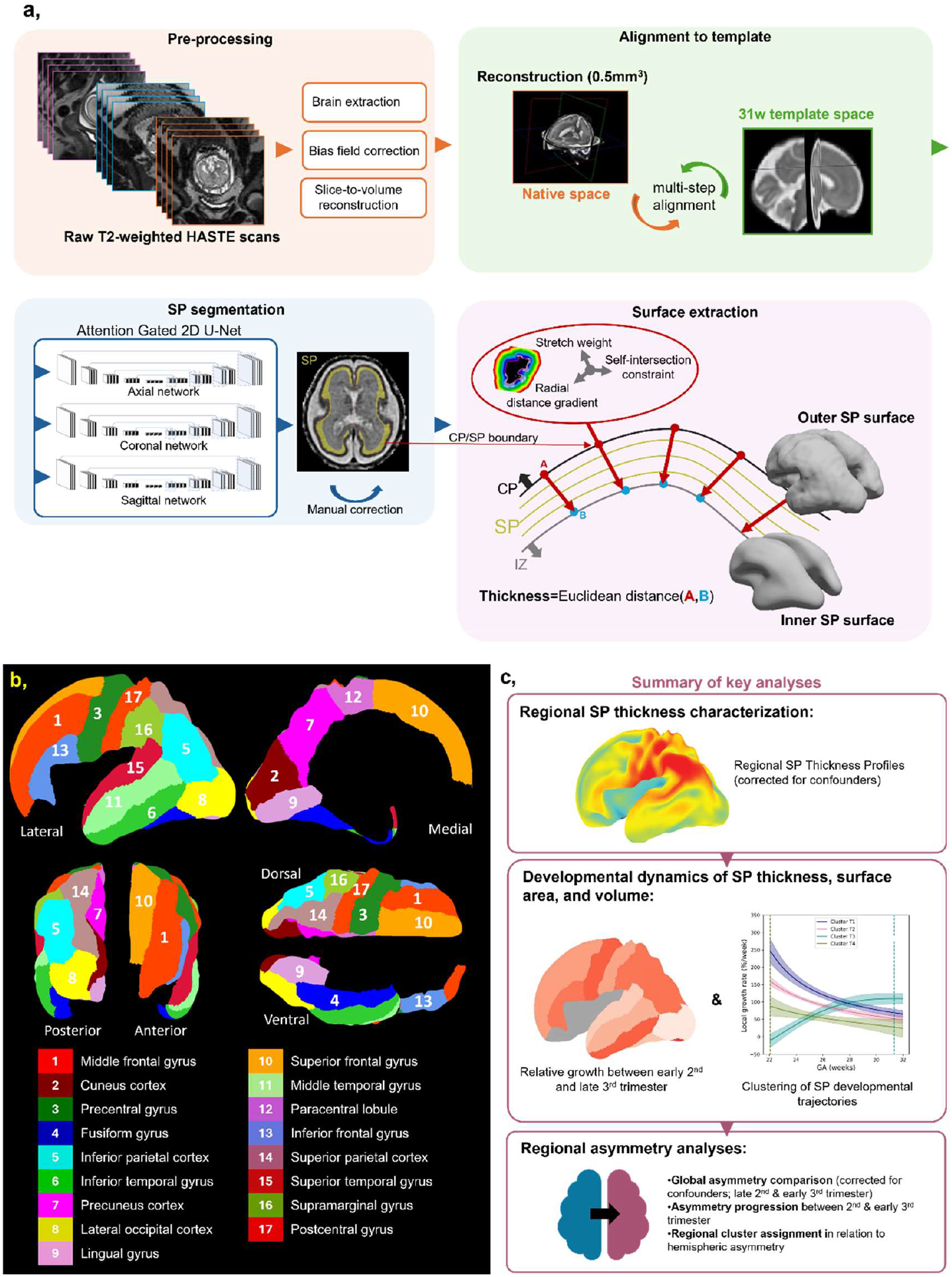
**a**, Schematic overview of the MRI processing pipeline. **b**, Visualization of the final set of regions used in the study, along with the corresponding color scheme. **c**, Summary of key analyses.

**Figure 2.**
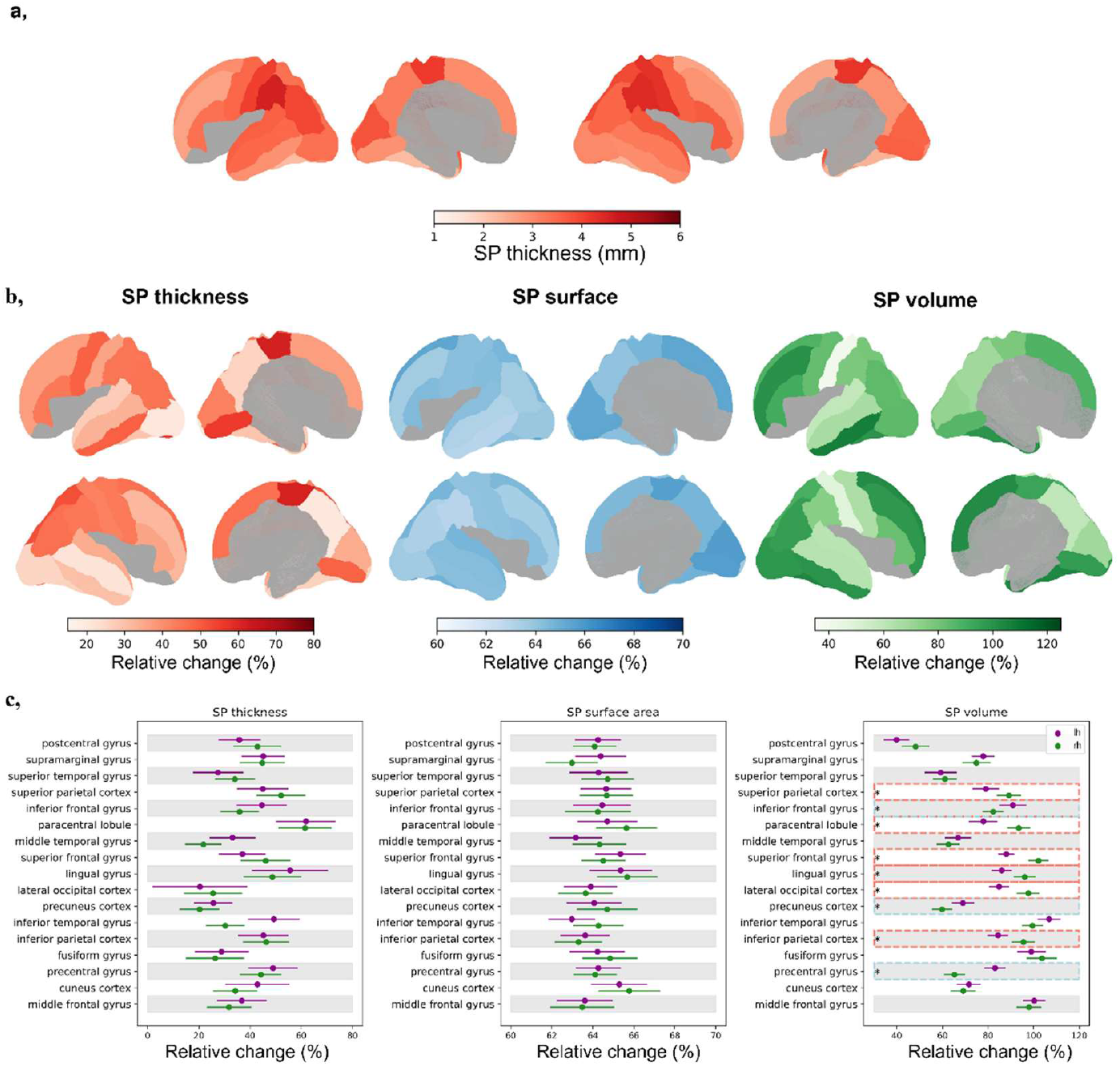
Regional variability of SP thickness and relative change of SP characteristics between late 2^nd^ and early 3^rd^ trimester. **a**, Regional variability in SP thickness after accounting for covariates: sulcal depth within subjects, and GA and residual brain volume within ROIs. **b**, Mean relative change in SP thickness, surface area, and volume projected onto SP surface maps, illustrating spatial patterns of growth. **c**, Regional distribution of relative growth, with dots indicating mean values and error bars representing confidence intervals. Regions outlined by boxes show significant left-right differences in relative growth across the period (FDR-corrected across regions within each metric) suggesting developmental asynchrony.

**Figure 3.**
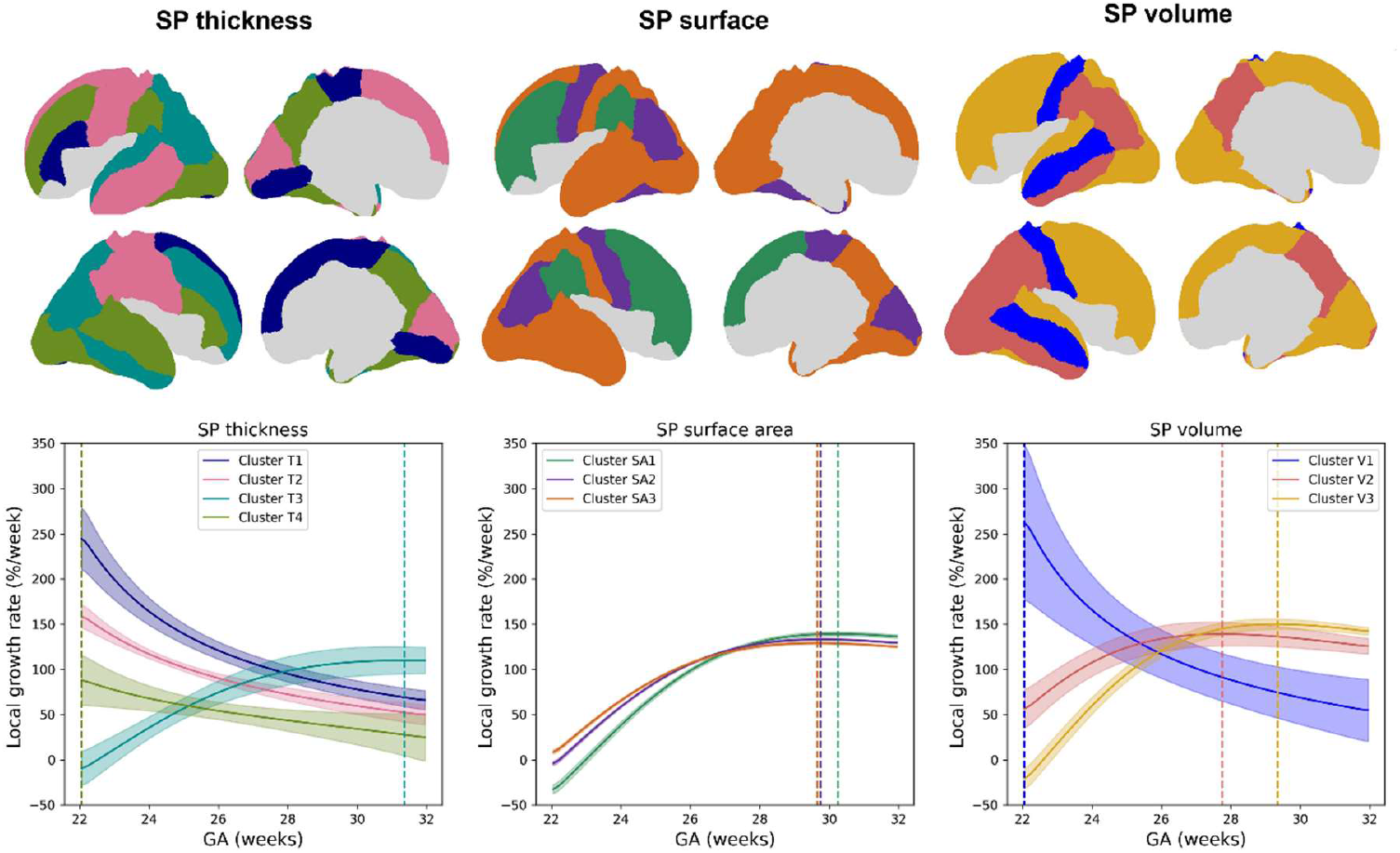
Clustering of SP developmental trajectories between 22–32 wGA. Top row: Results of hierarchical clustering (Ward’s method) applied to the regional developmental profiles for SP thickness, surface area, and volume. Bottom row: Cluster-averaged growth profiles showing mean local relative growth rates for each cluster over time and associated confidence intervals. Vertical lines indicate the time point of peak growth for each cluster.

**Figure 4.**
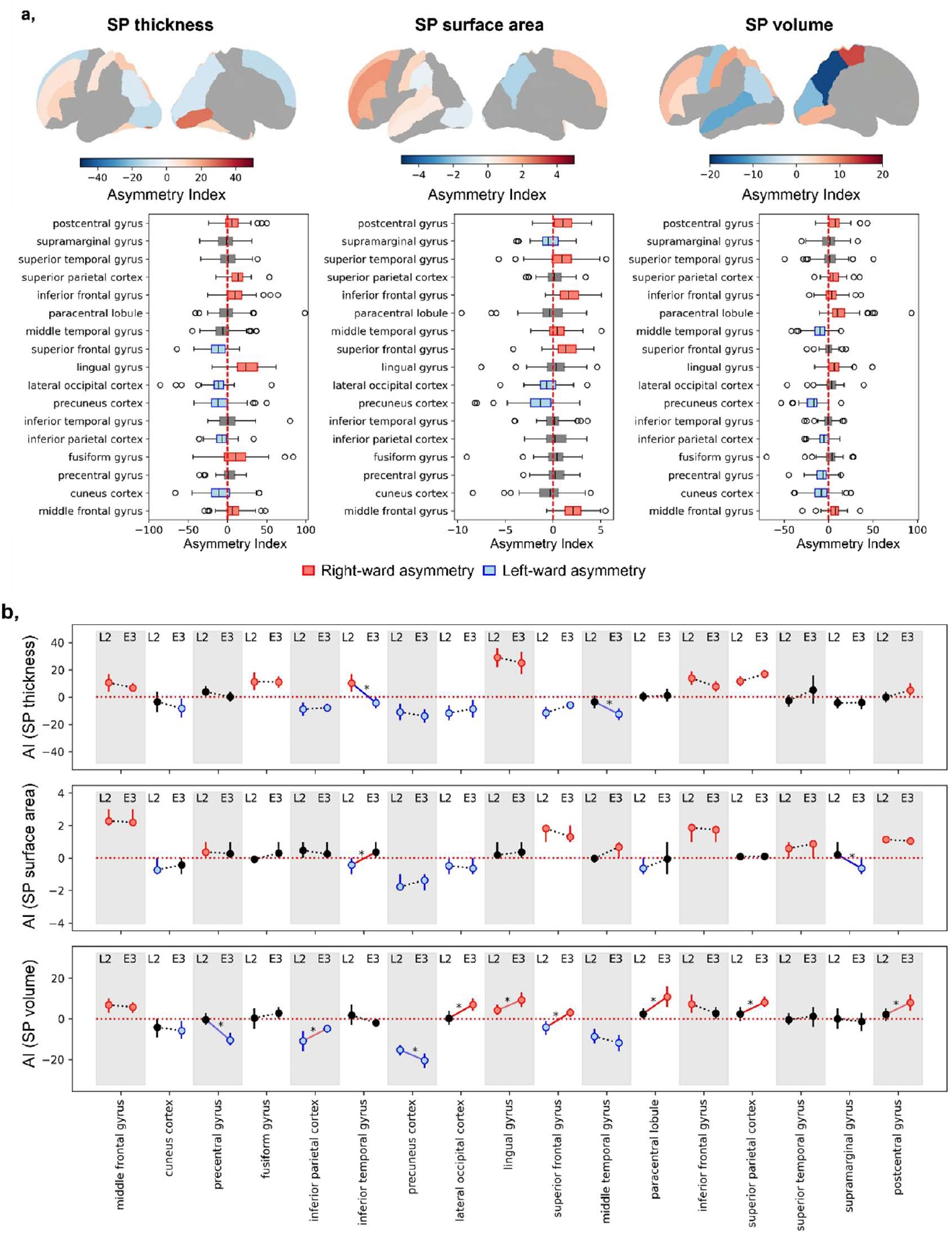
Asymmetry analysis. **a**, Asymmetry evaluations of regional SP characteristics: thickness, surface area, and volume. The upper row displays mean asymmetry index values projected onto cortical surface maps. The lower row presents the distribution of asymmetry indices across regions. Only regions exhibiting statistically significant asymmetries (p < 0.05, FDR-corrected across regions within metric) are shown in color on surface plots. Regions with significant rightward asymmetry are color-coded in red, while those with significant leftward asymmetry are shown in blue. **b**, Developmental changes in hemispheric asymmetry between late 2^nd^ and early 3^rd^ trimester. For each region, the asymmetry index (AI) is shown at late 2^nd^ (L2) and early 3^rd^ (E3) trimesters. Colored dots indicate significant lateralization (t-test vs. 0, FDR-corrected across regions within metric): red for rightward, blue for leftward, grey for non-significant. Dots represent mean AI; vertical lines show 95% confidence intervals. Lines connecting L2 and E3 indicate change in AI between the periods: red for rightward, blue for leftward shifts (Z-test on difference between timepoints, FDR-corrected across regions within metric), grey for non-significant changes.

### Extraction of whole-brain and regional SP characteristics

All measurements were computed in native space using registration parameters from the SP segmentation step. Whole-brain volume was the sum of CP, SP, and ‘other’ supratentorial labels multiplied by voxel volume. SP surface area was the total area of triangles on the bilateral *outer* SP surface, excluding the cingular pole. SP thickness was the Euclidean distance between corresponding inner and outer surface vertices, summarized as the median across hemispheres (excluding cingular pole). SP depth, measuring the cortical folding depth as amplitude of undulations of the outer SP surface, was computed similarly to sulcal depth (adapted from^103^ using MIRTK (v2.0) as the median displacement along surface normals from an inflated outer SP surface, rescaled so each subject’s minimum depth was zero across hemispheres (excluding cingular pole). Regional SP volume was voxel count within the region multiplied by voxel volume, surface area was summed triangle areas within the region, thickness and depth were medians of vertex-wise measurements within regions.

### Statistical analysis

Globally, our analyses first validated SP measurements and corrected for confounders (GA, residual brain volume, and regional SP depth), then assessed regional variability in SP thickness and interhemispheric asymmetry across SP characteristics. We next examined developmental changes in SP thickness, surface area, and volume from late 2^nd^ to early 3^rd^ trimester to identify the most developing regions and changes in asymmetry. Finally, we clustered regional growth trajectories to identify shared developmental patterns and evaluated whether lateralized features emerged within these profiles.

### Regional variation of SP characteristics and hemispheric asymmetries

To validate the extraction of SP characteristics, we first performed ANCOVA analysis to assess effects of GA, sex, and residual brain volume (corrected for GA) on whole-brain SP thickness, surface area, volume. Interaction terms (GA × sex, residual volume × sex) were tested. Subjects with unknown sex were coded as missing. Relationships were evaluated with F-statistics and associated p-values, and partial eta squared (ƞ^2^) to report effect sizes. Strong GA effects prompted post hoc visualization with linear or quadratic models chosen based on Akaike Information Criterion (AIC). Parameters were estimated with outlier-robust RANSAC regression^104^, with fixed seed (42) and minimum 75% sample inclusion. Note, this strategy was applied consistently across all analyses to residualize measures for covariate corrections (including the residual brain volume mentioned above). Because regional SP thickness could be confounded by geometric variation of outer SP surface undulations (for example, SP thickness tends to be thinner at the bottom of the SP grooves), a second ANCOVA tested thickness dependence on regional GA and residual brain volume as between-subject and SP depth (corrected for GA and residual brain volume) as within-region, within-hemisphere covariates. Significant within-subject region × depth interaction (Supp. Table 4) indicated that correcting for SP depth was necessary to isolate true regional thickness differences. Regional thickness values were thus pre-corrected in all analyses accordingly. Finally, we modelled regional variability in SP characteristics with GA and residual brain volume as between-subject covariates, and region and hemisphere as within-subject covariates to account for the nested data structure.

To isolate regional variation, metrics were then residualized for key covariates: GA and residual brain volume, and also for regional SP depth in case of thickness. Regional variation in SP volume and surface area was not interpreted due to parcellation dependence. Absolute left–right differences were assessed using paired t-tests with Benjamini–Hochberg FDR correction across regions for each SP characteristic. Additionally, normalized asymmetry indices (AI) were calculated as AI = [(right – left) × 100] / [0.5 × (right + left)], with positive values indicating rightward asymmetry. Significant lateralization was determined by one-sample t-tests against zero with FDR correction.

### Evolution of regional SP characteristics between late 2^nd^ and early 3^rd^ trimester

Subjects were grouped into late 2^nd^ and early 3^rd^ trimester groups based on GA at scan. Within each group, mean regional values were calculated after residualizing for GA and residual brain volume (and for thickness, also regional SP depth). The percent change in group means between the two periods was computed to normalize for baseline size, allowing cross-regional comparisons of SP surface area and volume. The same approach was applied to SP thickness for comparability, although absolute changes in thickness were also examined given their distinct interpretability. 95% confidence intervals (CIs) were derived from standard errors based on group standard deviations and sample sizes, assuming a normal distribution and a critical Z-value of 1.96. Developmental differences in relative growth between homotopic regions were assessed using Z-scores from percent change differences, with pooled standard errors calculated from the CIs. Asymmetry indices (AI) were computed within each group and tested for significance using one-sample t-tests with FDR correction. Changes in AI between groups were evaluated using Z-tests on differences in mean AI, with pooled standard errors, to identify regions showing age-dependent shifts in lateralization.

### Clustering to identify shared developmental dynamics

Regional SP values were corrected for residual brain volume (and for thickness, also for regional SP depth), then modelled against GA using linear or quadratic models selected by AIC within the RANSAC framework, balancing sensitivity to nonlinearities with resistance to outliers during this period of rapid growth prior to SP dissolution (~34–35wGA)^24^. More flexible models (e.g., splines) were avoided due to noise sensitivity. Continuous growth trajectories were sampled every 0.1 weeks between 22-32wGA to produce smooth developmental profiles. Instantaneous growth rates were estimated by finite differences, normalized by baseline value to express percent growth per week, allowing size-independent comparison across regions that emphasized developmental trajectory shapes rather than absolute magnitudes (this was also performed in case of thickness for consistency). Profiles were then clustered using hierarchical Ward’s method, which preserves nested structure and is robust across MRI-derived features^105^. to group regions with similar developmental dynamics, with the final number of clusters selected using silhouette scores and visual inspection of dendrograms. Resulting clusters showed relatively high internal consistency based on silhouette scores and low within-cluster variance (though this similarity might be influenced by the choice of the modelling strategy). Cluster-averaged profiles were used to characterize dominant patterns, calculating peak growth rate, its corresponding GA, the AUC as a measure of cumulative growth, and growth density, defined as the standard deviation of the smoothed relative growth rate curve to capture the temporal concentration of growth. Bilateral symmetry in clustering was assessed by calculating the proportion of homotopic region pairs assigned to the same cluster, normalized by the number of unique regions per cluster, with higher values indicating greater homotopic consistency in developmental trajectories.

## Acknowledgments

This work was supported by the National Institute of Biomedical Imaging and Bioengineering (R01EB031170) and the National Institute of Neurological Disorders and Stroke (K23NS101120, R01NS114087, and R01NS121334), National Institutes of Health. We thank Henry A. Feldman for his thoughtful feedback and insightful discussion regarding our analyses. We are deeply grateful to the families whose participation made this research possible.

## Data, materials, and software availability

Image processing code is available on GitHub (https://github.com/FNNDSC). Other processing used publicly available toolboxes detailed in *Materials and Methods*. Analyses were done mainly in Python (3.11.2) and RStudio (4.4.3). Manual corrections and some visualizations used Freeview (3.0). Anonymized data, including MRI scans and derivatives, will be shared upon reasonable request and signing a data usage agreement.

## References

1. Bystron, I., Blakemore, C. & Rakic, P. Development of the human cerebral cortex: Boulder Committee revisited. Nat Rev Neurosci 9, 110–122 (2008).

2. Gilmore, J. H., Knickmeyer, R. C. & Gao, W. Imaging structural and functional brain development in early childhood. Nat Rev Neurosci 19, 123–137 (2018).

3. Hüppi, P. S. Growth and Development of the Brain and Impact on Cognitive Outcomes. in 137–151 (2010). doi:10.1159/000281156.

4. Connors, S. L. et al. Fetal Mechanisms in Neurodevelopmental Disorders. Pediatr Neurol 38, 163–76 (2008).

5. Silbereis, J. C., Pochareddy, S., Zhu, Y., Li, M. & Sestan, N. The Cellular and Molecular Landscapes of the Developing Human Central Nervous System. Neuron 89, 248–268 (2016).

6. Vasung, L. et al. Quantitative and Qualitative Analysis of Transient Fetal Compartments during Prenatal Human Brain Development. Front Neuroanat 10, (2016).

7. Friauf, E., McConnell, S. & Shatz, C. Functional synaptic circuits in the subplate during fetal and early postnatal development of cat visual cortex. The Journal of Neuroscience 10, 2601–2613 (1990).

8. De Carlos, J. & O’Leary, D. Growth and targeting of subplate axons and establishment of major cortical pathways [published erratum appears in J Neurosci 1993 Mar;13(3):following table of contents]. The Journal of Neuroscience 12, 1194–1211 (1992).

9. Tolner, E. A., Sheikh, A., Yukin, A. Y., Kaila, K. & Kanold, P. O. Subplate Neurons Promote Spindle Bursts and Thalamocortical Patterning in the Neonatal Rat Somatosensory Cortex. The Journal of Neuroscience 32, 692–702 (2012).

10. Wess, J. M., Isaiah, A., Watkins, P. V. & Kanold, P. O. Subplate neurons are the first cortical neurons to respond to sensory stimuli. Proceedings of the National Academy of Sciences 114, 12602–12607 (2017).

11. Duque, A., Krsnik, Z., Kostović, I. & Rakic, P. Secondary expansion of the transient subplate zone in the developing cerebrum of human and nonhuman primates. Proceedings of the National Academy of Sciences 113, 9892–9897 (2016).

12. Kostovic, I., Judas, M., Rados, M. & Hrabac, P. Laminar Organization of the Human Fetal Cerebrum Revealed by Histochemical Markers and Magnetic Resonance Imaging. Cerebral Cortex 12, 536–544 (2002).

13. Krsnik, Ž., Majić, V., Vasung, L., Huang, H. & Kostović, I. Growth of Thalamocortical Fibers to the Somatosensory Cortex in the Human Fetal Brain. Front Neurosci 11, (2017).

14. Pogledic, I. et al. The Subplate Layers: The Superficial and Deep Subplate Can be Discriminated on 3 Tesla Human Fetal Postmortem MRI. Cerebral Cortex 30, 5038–5048 (2020).

15. Pogledic, I. et al. 3T MRI signal intensity profiles and thicknesses of transient zones in human fetal brain at mid-gestation. European Journal of Paediatric Neurology 35, 67–73 (2021).

16. Kostović, I. The enigmatic fetal subplate compartment forms an early tangential cortical nexus and provides the framework for construction of cortical connectivity. Prog Neurobiol 194, 101883 (2020).

17. Guo, X. et al. Molecular Lineages and Spatial Distributions of Subplate Neurons in the Human Fetal Cerebral Cortex. Advanced Science 11, (2024).

18. Kostović, I. Prenatal development of nucleus basalis complex and related fiber systems in man: A histochemical study. Neuroscience 17, 1047–1077 (1986).

19. Schwartz, M. L. & Goldman‐Rakic, P. S. Prenatal specification of callosal connections in rhesus monkey. Journal of Comparative Neurology 307, 144–162 (1991).

20. Verney, C. Distribution of the catecholaminergic neurons in the central nervous system of human embryos and fetuses. Microsc Res Tech 46, 24–47 (1999).

21. Kostovic, I. Laminar Organization of the Human Fetal Cerebrum Revealed by Histochemical Markers and Magnetic Resonance Imaging. Cerebral Cortex 12, 536–544 (2002).

22. Allendoerfer, K. L. & Shatz, C. J. The Subplate, A Transient Neocortical Structure: Its Role in the Development of Connections between Thalamus and Cortex. Annu Rev Neurosci 17, 185–218 (1994).

23. Luhmann, H. J., Kirischuk, S. & Kilb, W. The Superior Function of the Subplate in Early Neocortical Development. Front Neuroanat 12, (2018).

24. Kostović, I. & Judaš, M. The development of the subplate and thalamocortical connections in the human foetal brain. Acta Paediatr 99, 1119–1127 (2010).

25. Cadwell, C. R., Bhaduri, A., Mostajo-Radji, M. A., Keefe, M. G. & Nowakowski, T. J. Development and Arealization of the Cerebral Cortex. Neuron 103, 980–1004 (2019).

26. Kostović, I., Judaš, M. & Sedmak, G. Developmental history of the subplate zone, subplate neurons and interstitial white matter neurons: relevance for schizophrenia. International Journal of Developmental Neuroscience 29, 193–205 (2011).

27. Hadders-Algra, M. Early human brain development: Starring the subplate. Neurosci Biobehav Rev 92, 276–290 (2018).

28. Yamashita, Y. et al. MR imaging of the fetus by a HASTE sequence. American Journal of Roentgenology 168, 513–519 (1997).

29. Diogo, M. C. et al. Echo-planar FLAIR Sequence Improves Subplate Visualization in Fetal MRI of the Brain. Radiology 292, 159–169 (2019).

30. Kuklisova-Murgasova, M., Quaghebeur, G., Rutherford, M. A., Hajnal, J. V. & Schnabel, J. A. Reconstruction of fetal brain MRI with intensity matching and complete outlier removal. Med Image Anal 16, 1550–1564 (2012).

31. Ebner, M. et al. An automated framework for localization, segmentation and super-resolution reconstruction of fetal brain MRI. Neuroimage 206, 116324 (2020).

32. Xu, J. et al. NeSVoR: Implicit Neural Representation for Slice-to-Volume Reconstruction in MRI. IEEE Trans Med Imaging 42, 1707–1719 (2023).

33. You, S. et al. Automatic cortical surface parcellation in the fetal brain using attention-gated spherical U-net. Front Neurosci 18, (2024).

34. Perkins, L. et al. Exploring Cortical Subplate Evolution Using Magnetic Resonance Imaging of the Fetal Brain. Dev Neurosci 30, 211–220 (2008).

35. Rollins, C. K. et al. Regional Brain Growth Trajectories in Fetuses with Congenital Heart Disease. Ann Neurol 89, 143–157 (2021).

36. Scott, J. A. et al. Growth trajectories of the human fetal brain tissues estimated from 3D reconstructed in utero MRI. International Journal of Developmental Neuroscience 29, 529–536 (2011).

37. Sousa, H. S. et al. A Deep Learning Approach for Segmenting the Subplate and Proliferative Zones in Fetal Brain MRI. in 17–27 (2023). doi:10.1007/978-3-031-45544-5_2.

38. Wilson, S. et al. Dynamic changes in subplate and cortical plate microstructure at the onset of cortical folding in vivo. Preprint at 10.7554/eLife.100895.1 (2024).

39. Calixto, C. et al. Characterizing microstructural development in the fetal brain using diffusion MRI from 23 to 36 weeks of gestation. Cerebral Cortex 34, (2024).

40. Corbett-Detig, J. et al. 3D global and regional patterns of human fetal subplate growth determined in utero. Brain Struct Funct 215, 255–263 (2011).

41. Vasung, L. et al. Spatiotemporal Differences in the Regional Cortical Plate and Subplate Volume Growth during Fetal Development. Cerebral Cortex 30, 4438–4453 (2020).

42. Vasung, L. et al. Association between Quantitative MR Markers of Cortical Evolving Organization and Gene Expression during Human Prenatal Brain Development. Cerebral Cortex 31, 3610–3621 (2021).

43. Vasung, L. et al. Quantitative In vivo MRI Assessment of Structural Asymmetries and Sexual Dimorphism of Transient Fetal Compartments in the Human Brain. Cerebral Cortex 30, 1752–1767 (2020).

44. Hong, J. et al. Fetal Cortical Plate Segmentation Using Fully Convolutional Networks With Multiple Plane Aggregation. Front Neurosci 14, (2020).

45. Judaš, M., Sedmak, G. & Kostović, I. The significance of the subplate for evolution and developmental plasticity of the human brain. Front Hum Neurosci 7, (2013).

46. Sarnat, H. B. Transitory and Vestigial Structures of the Developing Human Nervous System. Pediatr Neurol 123, 86–101 (2021).

47. Zhang, Z. et al. Development of laminar organization of the fetal cerebrum at 3.0T and 7.0T: a postmortem MRI study. Neuroradiology 53, 177–184 (2011).

48. Kostović, I., Radoš, M., Kostović-Srzentić, M. & Krsnik, Ž. Fundamentals of the Development of Connectivity in the Human Fetal Brain in Late Gestation: From 24 Weeks Gestational Age to Term. J Neuropathol Exp Neurol 80, 393–414 (2021).

49. Gatti, M. G. et al. Functional maturation of neocortex: a base of viability. The Journal of Maternal-Fetal & Neonatal Medicine 25, 101–103 (2012).

50. Berretz, G., Wolf, O. T., Güntürkün, O. & Ocklenburg, S. Atypical lateralization in neurodevelopmental and psychiatric disorders: What is the role of stress? Cortex 125, 215–232 (2020).

51. Kong, X. et al. Mapping brain asymmetry in health and disease through the ENIGMA consortium. Hum Brain Mapp 43, 167–181 (2022).

52. Wang, B. et al. Brain asymmetry: a novel perspective on hemispheric network. Brain Science Advances 9, 56–77 (2023).

53. Kasprian, G. et al. The Prenatal Origin of Hemispheric Asymmetry: An In Utero Neuroimaging Study. Cerebral Cortex 21, 1076–1083 (2011).

54. Gilmore, J. H. et al. Regional Gray Matter Growth, Sexual Dimorphism, and Cerebral Asymmetry in the Neonatal Brain. The Journal of Neuroscience 27, 1255–1260 (2007).

55. Habas, P. A. et al. Early Folding Patterns and Asymmetries of the Normal Human Brain Detected from in Utero MRI. Cerebral Cortex 22, 13–25 (2012).

56. Vasung, L. et al. Quantitative In vivo MRI Assessment of Structural Asymmetries and Sexual Dimorphism of Transient Fetal Compartments in the Human Brain. Cerebral Cortex 30, 1752–1767 (2020).

57. Kienast, P. et al. The Prenatal Origins of Human Brain Asymmetry: Lessons Learned from a Cohort of Fetuses with Body Lateralization Defects. Cerebral Cortex 31, 3713–3722 (2021).

58. Taymourtash, A. et al. Fetal development of functional thalamocortical and cortico–cortical connectivity. Cerebral Cortex 33, 5613–5624 (2023).

59. Ouyang, M., Dubois, J., Yu, Q., Mukherjee, P. & Huang, H. Delineation of early brain development from fetuses to infants with diffusion MRI and beyond. Neuroimage 185, 836–850 (2019).

60. Gondova, A. et al. Attention-gated Convolutional Neural Network for Automated Segmentation of Fetal Subplate from MRI. in OHBM 2025 annual meeting (Brisbane, Australia, 2025).

61. Kyriakopoulou, V. et al. Normative biometry of the fetal brain using magnetic resonance imaging. Brain Struct Funct 222, 2295–2307 (2017).

62. Namburete, A. I. L. et al. Normative spatiotemporal fetal brain maturation with satisfactory development at 2 years. Nature 623, 106–114 (2023).

63. Radoš, M., Judaš, M. & Kostović, I. In vitro MRI of brain development. Eur J Radiol 57, 187–198 (2006).

64. Molliver, M. E., Kostović, I. & Van Der Loos, H. The development of synapses in cerebral cortex of the human fetus. Brain Res 50, 403–407 (1973).

65. Kostovic, I. & Rakic, P. Development of prestriate visual projections in the monkey and human fetal cerebrum revealed by transient cholinesterase staining. The Journal of Neuroscience 4, 25–42 (1984).

66. Kanold, P. O., Deng, R. & Meng, X. The Integrative Function of Silent Synapses on Subplate Neurons in Cortical Development and Dysfunction. Front Neuroanat 13, (2019).

67. Vasung, L. et al. Spatiotemporal Differences in the Regional Cortical Plate and Subplate Volume Growth during Fetal Development. Cerebral Cortex 30, 4438–4453 (2020).

68. McConnell, S. K., Ghosh, A. & Shatz, C. J. Subplate Neurons Pioneer the First Axon Pathway from the Cerebral Cortex. Science (1979) 245, 978–982 (1989).

69. Žunić Išasegi, I., Kopić, J., Smilović, D., Krsnik, Ž. & Kostović, I. Transient Subplate Sublayer Forms Unique Corridor for Differential Ingrowth of Associative Pulvinar and Primary Visual Projection in the Prospective Visual Cortical Areas of the Human Fetal Occipital Lobe. Cerebral Cortex 32, 110–122 (2021).

70. Rakic, P. Specification of Cerebral Cortical Areas. Science (1979) 241, 170–176 (1988).

71. Rubenstein, J. L. R. Genetic Control of Cortical Development. Cerebral Cortex 9, 521–523 (1999).

72. Kostović, I. & Judaš, M. Correlation between the sequential ingrowth of afferents and transient patterns of cortical lamination in preterm infants. Anat Rec 267, 1–6 (2002).

73. Kanold, P. O. & Luhmann, H. J. The Subplate and Early Cortical Circuits. Annu Rev Neurosci 33, 23–48 (2010).

74. Barber, M. & Pierani, A. Tangential migration of glutamatergic neurons and cortical patterning during development: Lessons from Cajal‐Retzius cells. Dev Neurobiol 76, 847–881 (2016).

75. Structural Development of the Human Prefrontal Cortex. in Handbook of Developmental Cognitive Neuroscience (The MIT Press, 2008). doi:10.7551/mitpress/7437.003.0017.

76. Hill, J. et al. Similar patterns of cortical expansion during human development and evolution. Proceedings of the National Academy of Sciences 107, 13135–13140 (2010).

77. Geschwind, D. H. & Rakic, P. Cortical Evolution: Judge the Brain by Its Cover. Neuron 80, 633–647 (2013).

78. Nishikuni, K. & Ribas, G. C. Study of fetal and postnatal morphological development of the brain sulci. J Neurosurg Pediatr 11, 1–11 (2013).

79. Bayly, P. V., Taber, L. A. & Kroenke, C. D. Mechanical forces in cerebral cortical folding: A review of measurements and models. J Mech Behav Biomed Mater 29, 568–581 (2014).

80. Budday, S. et al. Mechanical properties of gray and white matter brain tissue by indentation. J Mech Behav Biomed Mater 46, 318–330 (2015).

81. Rana, S. et al. The Subplate: A Potential Driver of Cortical Folding? Cerebral Cortex 29, 4697–4708 (2019).

82. Kostovic, I. & Rakic, P. Developmental history of the transient subplate zone in the visual and somatosensory cortex of the macaque monkey and human brain. J Comp Neurol 297, 441–470 (1990).

83. Margulies, D. S. et al. Situating the default-mode network along a principal gradient of macroscale cortical organization. Proceedings of the National Academy of Sciences 113, 12574–12579 (2016).

84. Dong, H.-M., Margulies, D. S., Zuo, X.-N. & Holmes, A. J. Shifting gradients of macroscale cortical organization mark the transition from childhood to adolescence. Proceedings of the National Academy of Sciences 118, (2021).

85. Huntenburg, J. M., Bazin, P.-L. & Margulies, D. S. Large-Scale Gradients in Human Cortical Organization. Trends Cogn Sci 22, 21–31 (2018).

86. Xu, J. et al. Tractography-based Parcellation of the Human Middle Temporal Gyrus. Sci Rep 5, 18883 (2015).

87. Kong, X.-Z. et al. Mapping cortical brain asymmetry in 17,141 healthy individuals worldwide via the ENIGMA Consortium. Proceedings of the National Academy of Sciences 115, (2018).

88. Sun, T. & Walsh, C. A. Molecular approaches to brain asymmetry and handedness. Nat Rev Neurosci 7, 655–662 (2006).

89. He, N., Palaniyappan, L., Linli, Z. & Guo, S. Abnormal hemispheric asymmetry of both brain function and structure in attention deficit/hyperactivity disorder: a meta-analysis of individual participant data. Brain Imaging Behav 16, 54–68 (2022).

90. Vanderauwera, J. et al. Atypical Structural Asymmetry of the Planum Temporale is Related to Family History of Dyslexia. Cerebral Cortex 28, 63–72 (2018).

91. Rollins, C. K. et al. Regional Brain Growth Trajectories in Fetuses with Congenital Heart Disease. Ann Neurol 89, 143–157 (2021).

92. Tustison, N. J. et al. N4ITK: Improved N3 Bias Correction. IEEE Trans Med Imaging 29, 1310–1320 (2010).

93. Jenkinson, M. Improved Optimization for the Robust and Accurate Linear Registration and Motion Correction of Brain Images. Neuroimage 17, 825–841 (2002).

94. Serag, A. et al. Construction of a consistent high-definition spatio-temporal atlas of the developing brain using adaptive kernel regression. Neuroimage 59, 2255–2265 (2012).

95. Lepage, C. et al. Human MR Evaluation of Cortical Thickness Using CIVET v2.1. in Organ Hum Brain Map. (2017).

96. Zhang, J. Fetal CP Surface Extraction ChRIS Plugin (2.1.1). Preprint at 10.5281/zenodo.17050400 (2025).

97. Taubin, G. Curve and surface smoothing without shrinkage. in Proceedings of IEEE International Conference on Computer Vision 852–857 (IEEE Comput. Soc. Press). doi:10.1109/ICCV.1995.466848.

98. Zhang, J. Self-Adapting Mesh Deformation for Fetal Brain Surface Fit (0.3.0). Preprint at 10.5281/zenodo.17050510 (2025).

99. Desikan, R. S. et al. An automated labeling system for subdividing the human cerebral cortex on MRI scans into gyral based regions of interest. Neuroimage 31, 968–980 (2006).

100. Yun, H. J. et al. Automatic labeling of cortical sulci for the human fetal brain based on spatio-temporal information of gyrification. Neuroimage 188, 473–482 (2019).

101. Robbins, S. Tuning and comparing spatial normalization methods. Med Image Anal 8, 311–323 (2004).

102. Boucher, M., Whitesides, S. & Evans, A. Depth potential function for folding pattern representation, registration and analysis. Med Image Anal 13, 203–214 (2009).

103. Makropoulos, A. et al. The developing human connectome project: A minimal processing pipeline for neonatal cortical surface reconstruction. Neuroimage 173, 88–112 (2018).

104. Fischler, M. A. & Bolles, R. C. Random sample consensus. Commun ACM 24, 381–395 (1981).

105. Thirion, B., Varoquaux, G., Dohmatob, E. & Poline, J.-B. Which fMRI clustering gives good brain parcellations? Front Neurosci 8, (2014).

