## Supplementary Materials for "Typical development of the human fetal subplate: regional heterogeneity, growth, and asymmetry assessed by in vivo T2-weighted MRI"

**This PDF file includes:**

Supp. Figures 1 to 8
Supp. Tables 1 to 8

**Figures**

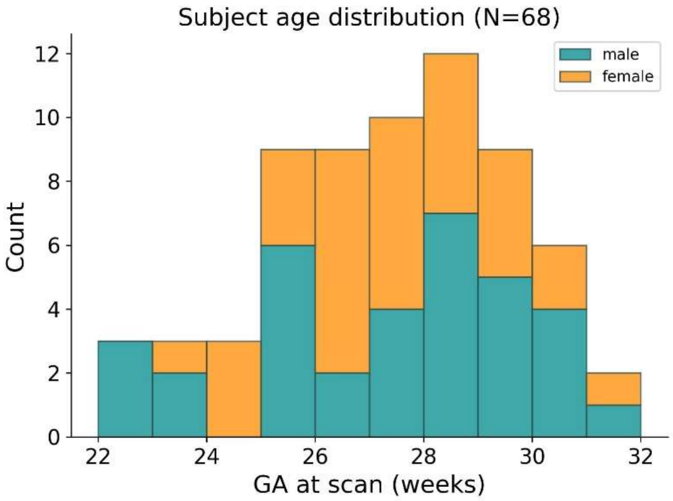


**Supp. Figure 1.** Age distribution across the cohort.

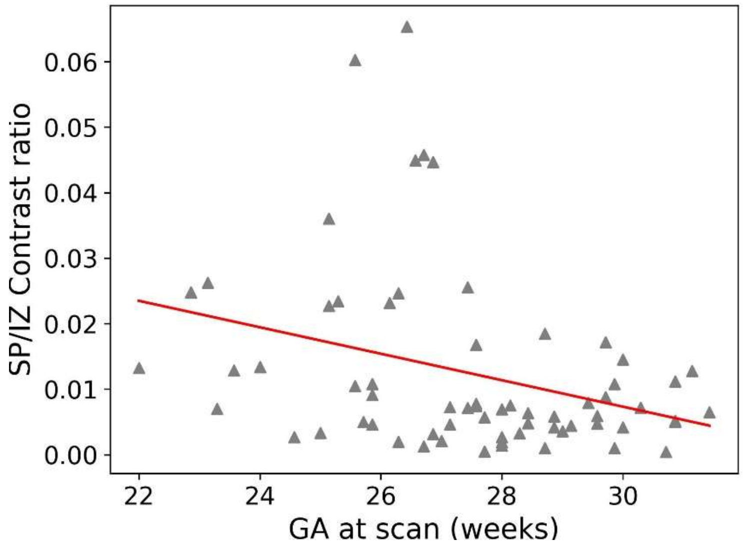


**Supp. Figure 2.** The contrast ratio of mean intensities at the boundary between SP and IZ (calculated as [mean SP signal - mean IZ signal] / [mean SP signal + mean IZ signal]) decreases with age ($R^2 = -0.334$, $p = 0.005$) making the two regions indistinguishable on T2-weighted MRI scans towards 32wGA.

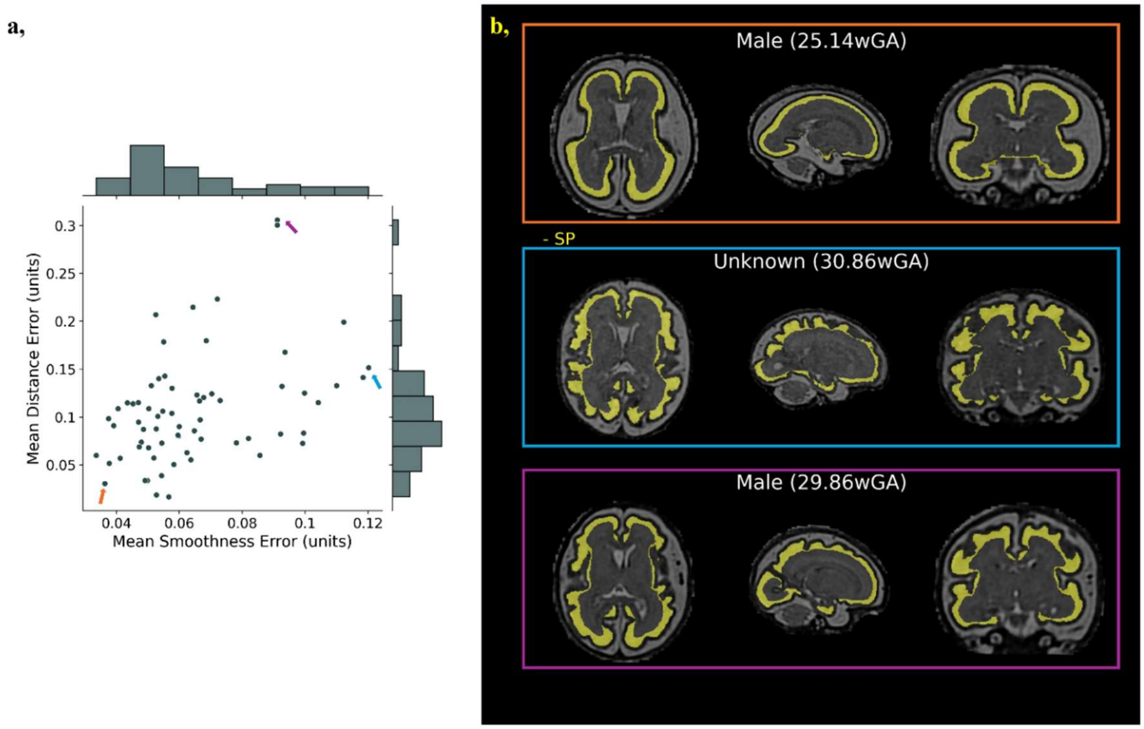


**Supp. Figure 3.** Surface extraction quality control (outer SP surface) across subjects and **b,** an example T2w reconstruction with overlayed SP segmentation (yellow) in the 31w template space for the subjects with the **best** and worst reconstruction quality (based on **smoothness** and **distance** errors).

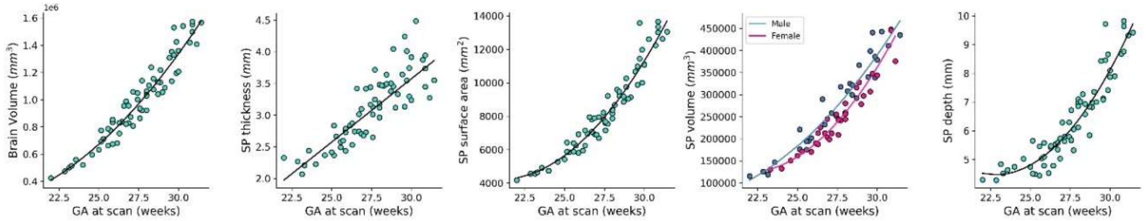


**Supp. Figure 4.** Relationships between gestational age (GA) and selected brain metrics across the cohort, presented as scatterplots. Each point represents a single subject. Metrics include SP thickness, surface area, thickness, depth, and whole brain volume (for reference). Sex is differentiated where significant sex-related effects or interactions with GA were identified in ANCOVA models (see *Supp. Table 3*): males are shown in blue, females in pink. Lines represent the best-fitting model (linear or quadratic), selected using Akaike Information Criterion (AIC), and are shown only where relationships were statistically significant ($p<0.05$).

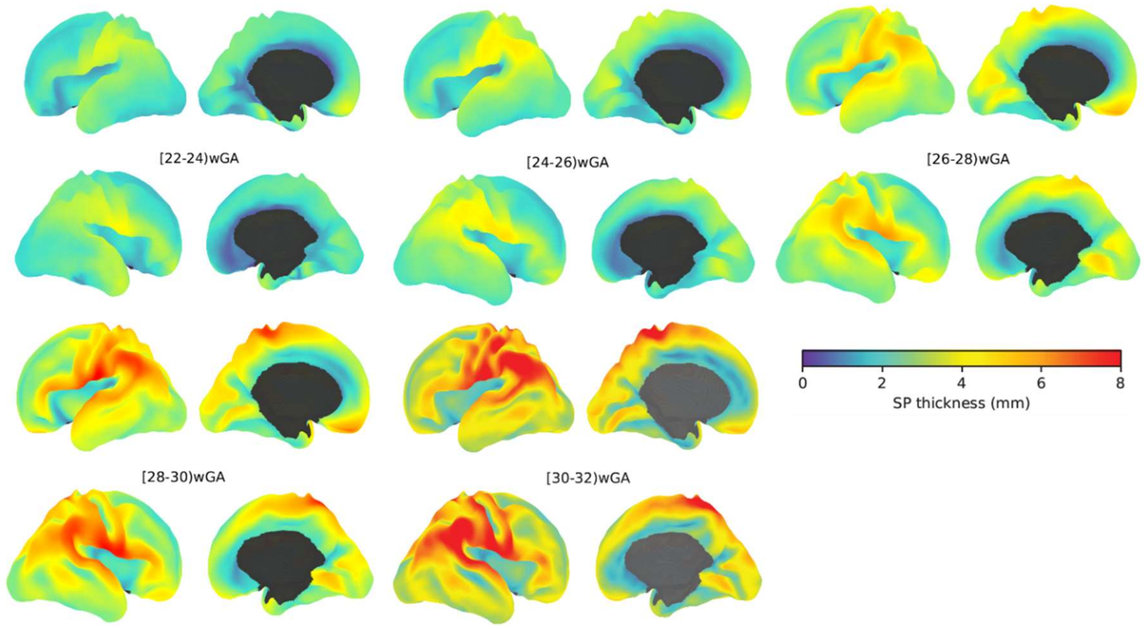


**Supp. Figure 5.** Example of SP thickness across gestation (22–32 weeks GA). Subjects were grouped into five gestational age bins, and mean vertex-wise SP thickness was computed within each bin for visualization.

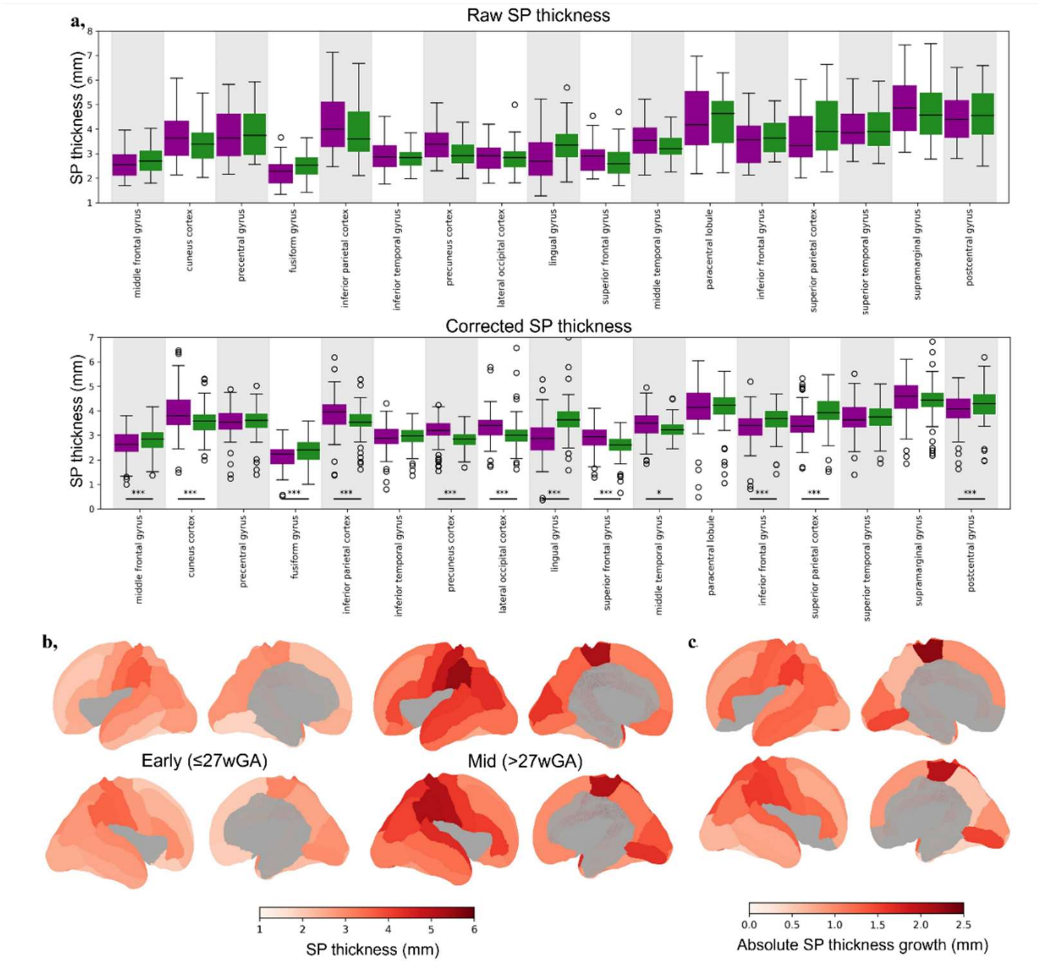


**Supp. Figure 6. a,** Boxplots summarizing raw regional SP thickness across subjects (top row) and values after correction for GA, residual brain volume, and regional SP sulcal depth (bottom row). After corrections, paired t-tests were used to assess differences between homotopic regions; statistically significant differences after FDR correction are marked with asterisks (* $p < 0.05$, ** $p < 0.01$, *** $p < 0.001$). Full numerical results, including paired comparisons for SP surface area and volume, are provided in *Supp. Table 6*. **b,** Regional SP thickness map comparing early and mid-gestational periods, corrected for covariates. **c,** Map of absolute regional SP thickness growth between early and mid-gestational periods.

**a,**

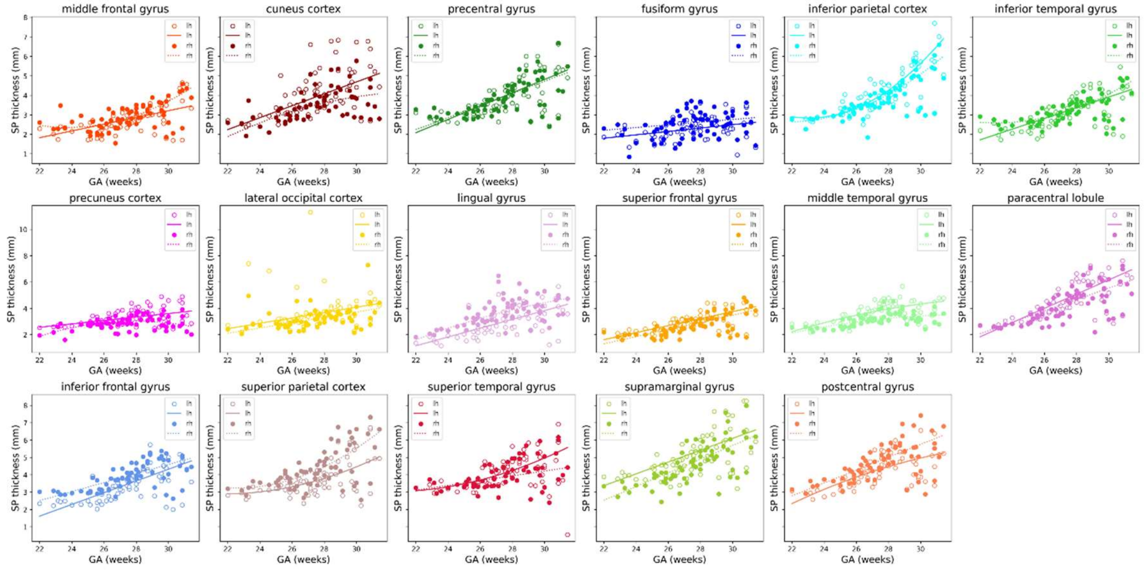


**b,**

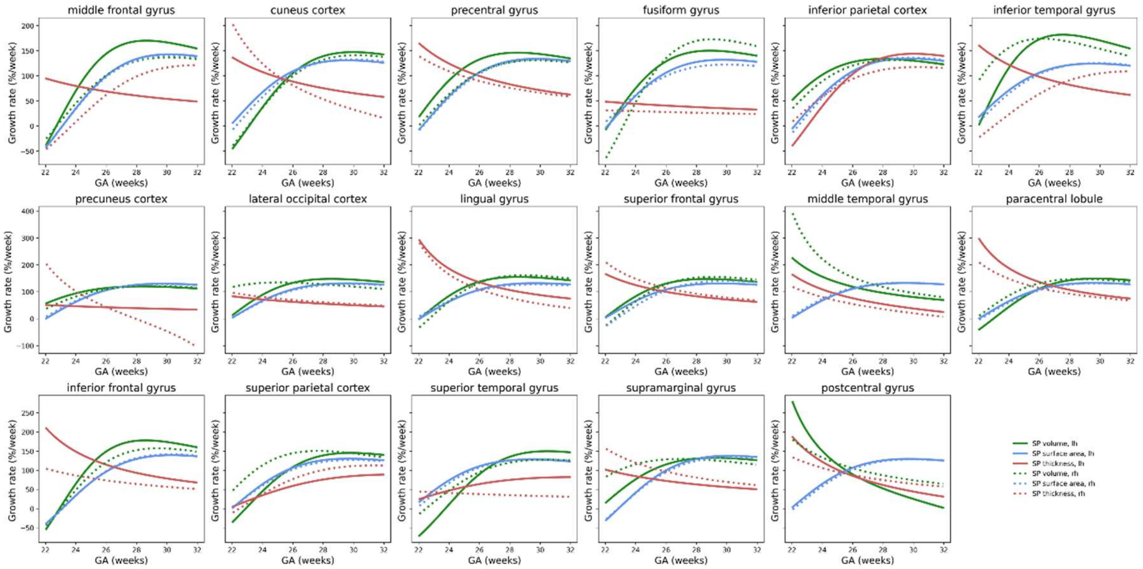


**Supp. Figure 7. a,** Developmental trajectories of regional SP thickness. Points represent regional thickness values after correction for confounders. The lines/curves show the RANSAC-derived developmental profiles. **b,** Regional relative growth rates (% per week) for SP characteristics sampled across 22–32wGA, based on RANSAC-fitted trajectories and high-resolution temporal sampling (0.1-week intervals). Instantaneous growth rates were computed using finite differences and normalized to account for regional size, yielding a consistent measure of developmental tempo across regions.

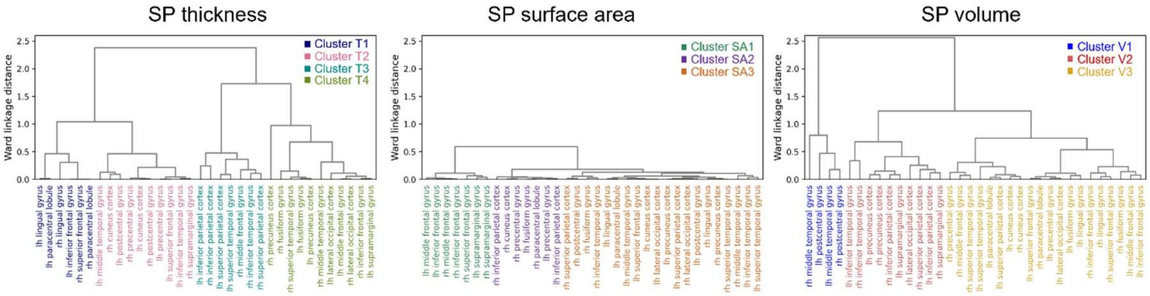


**Supp. Figure 8.** Dendrogram from hierarchical clustering (Ward's method) based on the developmental profiles shown in *Supp. Figure 7b*. The clustering reveals groups of SP regions with similar developmental trajectories over the studied gestational window.

### Tables

**Supp. Table 1.** Cohort characteristics. Reported p-values compare characteristics between early and mid-gestational periods and result from t-tests for continuous variables and χ² test for sex and maternal variables: categorized maternal BMI and education level.

| | TD (N=68) | Early (N=29) | Mid (N=39) | |
|---|---|---|---|---|
| **Males/Females/Unknown** | 34/32/2 | 13/14/1 | 21/17/1 | $\chi^2$= 0.212, p= 0.645 |
| **GA at scan (weeks) (median [range])** | 27.365 [22.00-31.43] | 25.57 [22.0-27.0] | 28.86 [27.14-31.43] | - |
| **Maternal age (years) (median [range])** | 32.565 [21.79-39.98] | 32.92 [22.71-39.90] | 32.94 [21.79-39.98] | t=-0.731, p=0.468 |
| **Maternal BMI (median [range])** | 26.584, [18.66-44.30] (N=61) | 26.58 [19.31-36.76] (N=26) | 32.94 [18.66-44.30] (N=35) | t=-0.020, p=0.984 |
| *Underweight* | 0 | 0 | 0 | $\chi^2$= 0.441, p= 0.802 |
| *Healthy Weigh* | 28 | 11 | 14 | |
| *Overweight* | 19 | 8 | 10 | |
| *Obesity* | 14 | 7 | 5 | |
| **Education** | | | | |
| *< College* | 1 | 0 | 1 | $\chi^2$= 1.156, p= 0.764 |
| *Partial College/ Specialized training/ High School* | 14 | 6 | 8 | |
| *Graduated standard College/University* | 21 | 8 | 13 | |
| *Graduate Degree* | 32 | 15 | 17 | |

**Supp. Table 2.** 21-label parcellation scheme compared to original DKT cortical regions.

| DKT label names | 21 label | Refered name |
|---|---|---|
| caudal middle frontal gyrus, rostral middle frontal gyrus | 1 | middle frontal gyrus |
| cuneus cortex, pericalcarine cortex (upper bank) | 2 | cuneus cortex |
| precentral gyrus | 3 | precentral gyrus |
| fusiform gyrus | 4 | fusiform gyrus |
| inferior parietal lobule | 5 | inferior parietal cortex |
| inferior temporal gyrus, temporal pole | 6 | inferior temporal gyrus |
| precuneus | 7 | precuneus cortex |
| lateral occipical cortex | 8 | lateral occipical cortex |
| lingual gurys, pericalcarine cortex (lower bank) | 9 | lingual gyrus |
| superior frontal gyrus | 10 | superior frontal gyrus |
| middle temporal gyrus, superior temporal sulcus, temporal pole | 11 | middle temporal gyrus |
| paracentral lobule | 12 | paracentral lobule |
| pars opercularis, pars orbitalis, pars triangularis | 13 | inferior frontal gyrus |
| superior parietal cortex | 14 | superior parietal cortex |
| superior temporal gyrus, superior temporal sulcus, transverse temporal gyrus, temporal pole | 15 | superior temporal gyrus |
| supramarginal gyrus | 16 | supramarginal gyrus |
| postcentral gyrus | 17 | postcentral gyrus |
| lateral orbitofrontal gyrus, medial orbitofrontal gyrus, frontal pole | 18 | orbitofrontal gyrus |
| insula | 19 | insula |
| caudal anterior cingulate cortex, posterior cingulate cortex, rostral anterior cingulate cortex, isthmus cingulate cortex | 20 | cingulate cortex |
| parahippocampal gyrus, entorhinal cortex | 21 | parahippocampal cortex |

**Supp. Table 3.** Results of ANCOVA analysis examining the relationship between the SP characteristics and covariates, including GA, residual brain volume (corrected for GA), and sex, along with relevant interactions. Legend: F: F-statistic from ANCOVA; PR (>F): statistical significance; $\eta^2$ = effect size.

| | | ***SP thickness*** | | | ***SP surface area*** | | | ***SP volume*** | | | ***SP depth*** | | |
|---|---|---|---|---|---|---|---|---|---|---|---|---|---|
| | | **F** | **PR (>F)** | **η²** | **F** | **PR (>F)** | **η²** | **F** | **PR (>F)** | **η²** | **F** | **PR (>F)** | **η²** |
| ***Continuous*** | | | | | | | | | | | | | |
| | GA | 393.553 | <0.001 | 0.774 | 6167.471 | <0.001 | 0.950 | 2823.253 | <0.001 | 0.927 | 380.265 | <0.001 | 0.850 |
| | residual brain volume | 52.507 | <0.001 | 0.103 | 248.569 | <0.001 | 0.038 | 156.999 | <0.001 | 0.052 | 5.542 | 0.022 | 0.012 |
| ***Categorical*** | | | | | | | | | | | | | |
| | Sex | 1.837 | 0.180 | 0.004 | 1.132 | 0.292 | 0.000 | 5.594 | 0.021 | 0.002 | 0.006 | 0.939 | 0.000 |
| ***Interactions*** | | | | | | | | | | | | | |
| | GA : sex | 0.263 | 0.610 | 0.001 | 13.120 | 0.001 | 0.002 | 1.059 | 0.308 | 0.000 | 1.650 | 0.204 | 0.004 |
| | residual brain volume : sex | 0.220 | 0.641 | 0.000 | 0.000 | 0.993 | 0.000 | 0.000 | 0.988 | 0.000 | 0.086 | 0.770 | 0.000 |

**Supp. Table 4.** Results of ANCOVA examining the relationship between the log-transformed regional SP thickness with SP depth (corrected for GA and residual brain volume), and across regions. The models account for hierarchical structure of the repeated measures across regions and hemispheres with a nested error term. See *Supp. Table 3* for legend.

| | | ***SP thickness*** | | |
|---|---|---|---|---|
| | | **F** | **PR (>F)** | **$\eta^2$** |
| ***Between*** | | | | |
| | GA | 920.094 | <0.001 | 0.383 |
| | residual brain volume | 66.957 | <0.001 | 0.028 |
| | SP depth | 20.229 | <0.001 | 0.008 |
| | SP depth : region | 2.287 | 0.014 | 0.015 |
| ***Subject:ROI*** | | | | |
| | SP depth | 190.129 | <0.001 | 0.016 |
| | ROI | 282.705 | <0.001 | 0.383 |
| | SP depth : region | 2.321 | 0.002 | 0.003 |
| ***Subject:ROI:hemi*** | | | | |
| | hemi | 3.813 | 0.051 | 0.000 |
| | SP depth | 295.686 | <0.001 | 0.010 |
| | SP depth : region | 6.329 | <0.001 | 0.004 |

**Supp. Table 5.** Results of ANCOVA analysis of regional differences and regional hemispheric asymmetries. The models account for hierarchical structure of the repeated measures across regions and hemispheres with a nested error term. See *Supp. Table 3* for legend.

| | | ***SP thickness**** | | | ***SP surface area*** | | | ***SP volume*** | | |
|---|---|---|---|---|---|---|---|---|---|---|
| | | **F** | **PR (>F)** | **η²** | **F** | **PR (>F)** | **η²** | **F** | **PR (>F)** | **η²** |
| ***Between*** | | | | | | | | | | |
| | GA | 94.698 | <0.001 | 0.260 | 5519.100 | <0.001 | 0.422 | 2371.900 | <0.001 | 0.423 |
| | residual brain volume | 4.309 | 0.042 | 0.012 | 224.100 | <0.001 | 0.017 | 141.900 | <0.001 | 0.025 |
| ***Subject:ROI*** | | | | | | | | | | |
| | region | 111.956 | <0.001 | 0.271 | 36981.000 | <0.001 | 0.554 | 1260.640 | <0.001 | 0.492 |
| | GA : region | 9.052 | <0.001 | 0.022 | 5.550 | <0.001 | 0.000 | 23.520 | <0.001 | 0.009 |
| ***Subject:ROI:hemi*** | | | | | | | | | | |
| | hemi | 8.945 | 0.003 | 0.001 | 70.700 | <0.001 | 0.000 | 0.744 | 0.389 | 0.000 |
| | GA : hemi | 2.347 | 0.126 | 0.000 | 13.660 | 0.001 | 0.000 | 2.241 | 0.135 | 0.000 |
| | region : hemi | 26.278 | <0.001 | 0.025 | 26.780 | <0.001 | 0.000 | 29.731 | <0.001 | 0.004 |
| | GA: region : hemi | 4.215 | <0.001 | 0.004 | 3.180 | <0.001 | 0.000 | 6.808 | <0.001 | 0.001 |

*SP thickness: residual thickness corrected for regional SP depth within subjects.

**Supp. Table 6.** Asymmetry analyses. The table reports t-statistics (T) and Cohen's D values derived from one-sample t-tests comparing asymmetry indices to zero to assess significant lateralization. p(corr) denotes FDR-corrected p-values, adjusted for multiple comparisons across regions within each metric.

| | | ***SP volume**** | | | | ***SP surface**** | | | | ***SP thickness**** | | | |
|---|---|---|---|---|---|---|---|---|---|---|---|---|---|
| RH-ward | **ROI** | **AI** | **T** | **Cohen's D** | **p (corr.)** | **AI** | **T** | **Cohen's D** | **p (corr.)** | **AI** | **T** | **Cohen's D** | **p (corr.)** |
| LH-ward | middle frontal gyrus | 5.95 | 5.73 | 0.61 | <0.001 | 2.23 | 13.31 | 0.57 | <0.001 | 5.74 | 3.19 | 0.28 | <0.001 |
| | cuneus cortex | -8.61 | -5.02 | 0.68 | <0.001 | -0.52 | -2.18 | 0.15 | 0.053 | -6.36 | -3.30 | 0.39 | <0.001 |
| | precentral gyrus | -7.75 | -6.72 | 0.67 | <0.001 | 0.31 | 2.16 | 0.11 | 0.053 | 1.59 | 1.11 | 0.07 | 0.364 |
| | fusiform gyrus | 1.82 | 2.14 | 0.16 | 0.047 | 0.18 | 0.89 | 0.06 | 0.441 | 5.86 | 3.46 | 0.29 | <0.001 |
| | inferior parietal cortex | -5.48 | -5.63 | 0.59 | <0.001 | 0.32 | 2.00 | 0.11 | 0.070 | -5.85 | -5.10 | 0.33 | <0.001 |
| | inferior temporal gyrus | -0.60 | -0.70 | 0.07 | 0.588 | 0.11 | 0.74 | 0.04 | 0.480 | 2.16 | 0.49 | 0.04 | 0.726 |
| | precuneus cortex | -18.11 | -13.02 | 1.59 | <0.001 | -1.49 | -6.08 | 0.42 | <0.001 | -10.57 | -6.20 | 0.74 | <0.001 |
| | lateral occipital cortex | 2.28 | 2.30 | 0.22 | 0.035 | -0.58 | -3.39 | 0.17 | 0.003 | -12.16 | -5.39 | 0.58 | <0.001 |
| | lingual gyrus | 6.80 | 4.65 | 0.60 | <0.001 | 0.32 | 1.34 | 0.09 | 0.244 | 34.59 | 11.06 | 1.31 | <0.001 |
| | superior frontal gyrus | -0.01 | 0.10 | 0.01 | 0.950 | 1.47 | 8.35 | 0.48 | <0.001 | -7.11 | -5.51 | 0.34 | <0.001 |
| | middle temporal gyrus | -10.43 | -7.38 | 0.77 | <0.001 | 0.46 | 2.75 | 0.14 | 0.017 | -3.95 | -2.94 | 0.30 | 0.011 |
| | paracentral lobule | 13.24 | 7.15 | 0.90 | <0.001 | -0.23 | -0.87 | 0.07 | 0.441 | 1.63 | -0.01 | 0.00 | 0.367 |
| | inferior frontal gyrus | 4.00 | 2.82 | 0.32 | 0.010 | 1.77 | 10.84 | 0.43 | <0.001 | 10.48 | 5.70 | 0.47 | <0.001 |
| | superior parietal cortex | 6.35 | 6.05 | 0.46 | <0.001 | 0.11 | 0.71 | 0.03 | 0.480 | 14.22 | 11.77 | 0.67 | <0.001 |
| | superior temporal gyrus | 0.68 | 0.10 | 0.01 | 0.950 | 0.78 | 3.43 | 0.23 | 0.003 | 6.51 | 2.76 | 0.28 | 0.353 |
| | supramarginal gyrus | 0.00 | 0.06 | 0.01 | 0.950 | -0.38 | -2.44 | 0.11 | 0.033 | -1.10 | -0.87 | 0.07 | 0.229 |
| | postcentral gyrus | 7.86 | 6.10 | 0.59 | <0.001 | 1.07 | 7.30 | 0.37 | <0.001 | 7.02 | 4.37 | 0.38 | 0.001 |

* SP volume / SP surface: corrected for GA and residual brain volume; SP thickness additionally corrected for regional SP depth within subject.

**Supp. Table 7.** Differences in relative growth and asymmetry between early and mid-gestational periods. p(corr) denotes FDR-corrected p-values, adjusted for multiple comparisons across regions within each metric. The Left/Right growth columns summarize mean regional values of each SP characteristic at the early and mid-gestational periods. % Growth indicates the relative percent increase over this interval, with corresponding 95% confidence intervals (CI). Left vs. Right columns compare relative percent growth between hemispheres using a Z-test based on the difference in percent change, accounting for propagated standard error. For Asymmetry Indices (AI), values are reported for the early and mid-gestational periods. The adjacent p(corr) columns show FDR-corrected p-values from one-sample t-tests against zero, corrected across regions. ΔAI reflects the absolute change in asymmetry between the two periods, with 95% CI. These changes were also compared using a Z-test that accounts for propagated standard error.

| | early ≤ 27wGA, mid > 27wGA | *SP thickness** | | | | | | | | | | | | | | | | | |
|---|---|---|---|---|---|---|---|---|---|---|---|---|---|---|---|---|---|---|---|
| | | left growth | | | | right growth | | | | left vs right | | AI (%) | | | | | | | |
| RH-ward | ROI | early | mid | %growth | CI | early | mid | %growth | CI | Z | p (corr.) | early AI | p (corr.) | mid AI | p (corr.) | ΔAI | CI [AI/100] | Z | p (corr.) |
| LH-ward | middle frontal gyrus | 2.15 | 2.94 | 36.84 | [27.06, 46.62] | 2.38 | 3.14 | 31.80 | [23.07, 40.52] | 0.75 | 0.665 | 10.57 | 0.005 | 6.89 | 0.000 | 3.67 | [-0.03, 0.11] | 1.03 | 0.537 |
| | cuneus cortex | 3.06 | 4.37 | 42.88 | [30.47, 55.3] | 2.92 | 3.93 | 34.27 | [25.58, 42.97] | 1.11 | 0.646 | -3.37 | 0.396 | -8.31 | 0.037 | 4.94 | [-0.05, 0.15] | 1.00 | 0.537 |
| | precentral gyrus | 2.77 | 4.13 | 49.00 | [39.42, 58.58] | 2.87 | 4.13 | 44.21 | [36.07, 52.34] | 0.75 | 0.665 | 3.95 | 0.070 | 0.30 | 0.868 | 3.65 | [-0.01, 0.09] | 1.40 | 0.362 |
| | fusiform gyrus | 1.88 | 2.43 | 28.98 | [18.44, 39.53] | 2.14 | 2.70 | 26.39 | [15.08, 37.7] | 0.33 | 0.902 | 11.29 | 0.004 | 11.09 | 0.000 | 0.20 | [-0.07, 0.08] | 0.05 | 0.959 |
| | inferior parietal cortex | 3.12 | 4.53 | 45.24 | [35.29, 55.18] | 2.85 | 4.17 | 46.30 | [37.33, 55.28] | -0.16 | 0.964 | -8.84 | 0.003 | -7.79 | 0.000 | 1.05 | [-0.04, 0.06] | 0.39 | 0.845 |
| | inferior temporal gyrus | 2.32 | 3.46 | 49.26 | [39.26, 59.25] | 2.54 | 3.31 | 30.41 | [22.79, 38.03] | 2.94 | 0.056 | 10.27 | 0.006 | -4.19 | 0.048 | 14.47 | [0.07, 0.22] | 3.93 | 0.001 |
| | precuneus cortex | 2.83 | 3.55 | 25.70 | [18.08, 33.33] | 2.55 | 3.07 | 20.40 | [12.57, 28.24] | 0.95 | 0.646 | -10.92 | 0.001 | -13.88 | 0.000 | 2.96 | [-0.04, 0.1] | 0.80 | 0.597 |
| | lateral occipital cortex | 3.04 | 3.66 | 20.45 | [2.04, 38.86] | 2.62 | 3.29 | 25.56 | [14.17, 36.95] | -0.46 | 0.842 | -11.70 | 0.000 | -8.60 | 0.022 | 3.09 | [-0.05, 0.11] | 0.74 | 0.597 |
| | lingual gyrus | 2.18 | 3.39 | 55.64 | [40.71, 70.57] | 2.89 | 4.29 | 48.73 | [37.43, 60.03] | 0.72 | 0.665 | 29.22 | 0.000 | 25.14 | 0.000 | 4.09 | [-0.06, 0.14] | 0.78 | 0.597 |
| | superior frontal gyrus | 2.38 | 3.26 | 37.09 | [27.97, 46.2] | 2.12 | 3.09 | 46.10 | [36.41, 55.79] | -1.33 | 0.646 | -11.45 | 0.000 | -5.80 | 0.001 | 5.66 | [0.01, 0.11] | 2.24 | 0.107 |
| | middle temporal gyrus | 2.96 | 3.94 | 33.16 | [24.26, 42.05] | 2.84 | 3.46 | 21.86 | [14.77, 28.94] | 1.95 | 0.437 | -3.40 | 0.151 | -12.45 | 0.000 | 9.05 | [0.03, 0.15] | 2.95 | 0.027 |
| | paracentral lobule | 3.12 | 5.04 | 61.81 | [50.2, 73.43] | 3.12 | 5.04 | 61.46 | [51.14, 71.78] | 0.05 | 0.964 | 0.53 | 0.819 | 1.13 | 0.663 | 0.59 | [-0.05, 0.06] | 0.21 | 0.948 |
| | inferior frontal gyrus | 2.68 | 3.88 | 44.62 | [34.83, 54.41] | 3.06 | 4.16 | 35.93 | [28.41, 43.45] | 1.38 | 0.646 | 13.85 | 0.000 | 7.68 | 0.001 | 6.17 | [0, 0.12] | 2.03 | 0.143 |
| | superior parietal cortex | 2.74 | 3.97 | 44.98 | [34.98, 54.98] | 3.08 | 4.69 | 52.08 | [42.5, 61.67] | -1.00 | 0.646 | 11.53 | 0.000 | 17.05 | 0.000 | 5.52 | [0.01, 0.1] | 2.41 | 0.090 |
| | superior temporal gyrus | 3.22 | 4.10 | 27.54 | [17.72, 37.36] | 3.14 | 4.21 | 34.11 | [26.24, 41.98] | -1.02 | 0.646 | -2.45 | 0.352 | 5.21 | 0.365 | 7.66 | [-0.03, 0.19] | 1.37 | 0.362 |
| | supramarginal gyrus | 3.72 | 5.39 | 45.12 | [36.67, 53.58] | 3.57 | 5.18 | 44.80 | [36, 53.59] | 0.05 | 0.964 | -4.24 | 0.070 | -4.01 | 0.100 | 0.23 | [-0.06, 0.06] | 0.08 | 0.959 |
| | postcentral gyrus | 3.46 | 4.70 | 35.91 | [27.72, 44.1] | 3.48 | 4.97 | 42.86 | [33.62, 52.1] | -1.10 | 0.646 | -0.01 | 0.996 | 5.13 | 0.037 | 5.14 | [-0.01, 0.11] | 1.71 | 0.247 |

| | *SP surface** | | | | | | | | | | | | | | | | | |
|---|---|---|---|---|---|---|---|---|---|---|---|---|---|---|---|---|---|---|
| | left growth | | | | right growth | | | | left vs right | | AI (%) | | | | | | | |
| ROI | early | mid | %growth | CI | early | mid | %growth | CI | Z | p (corr.) | early AI | p (corr.) | mid AI | p (corr.) | ΔAI | CI [AI/100] | Z | p (corr.) |
| middle frontal gyrus | 235.42 | 385.16 | 63.61 | [62.25, 64.96] | 235.42 | 393.80 | 67.28 | [61.93, 65.04] | 0.12 | 0.962 | 2.28 | 0.000 | 2.20 | 0.000 | 0.08 | [-0.005, 0.007] | 0.26 | 0.842 |
| cuneus cortex | 91.67 | 151.51 | 65.29 | [63.92, 66.65] | 91.67 | 150.87 | 64.59 | [64.28, 67.28] | -0.47 | 0.960 | -0.73 | 0.015 | -0.43 | 0.263 | 0.31 | [-0.005, 0.011] | 0.76 | 0.786 |
| precentral gyrus | 195.97 | 321.92 | 64.27 | [63.17, 65.37] | 195.97 | 322.80 | 64.72 | [63.07, 65.17] | 0.19 | 0.960 | 0.37 | 0.039 | 0.28 | 0.185 | 0.09 | [-0.004, 0.005] | 0.39 | 0.786 |
| fusiform gyrus | 106.05 | 174.17 | 64.22 | [62.88, 65.57] | 106.05 | 174.68 | 64.71 | [63.51, 66.17] | -0.64 | 0.960 | -0.08 | 0.721 | 0.30 | 0.311 | 0.38 | [-0.003, 0.01] | 1.17 | 0.688 |
| inferior parietal cortex | 157.19 | 257.21 | 63.63 | [62.43, 64.83] | 157.19 | 257.86 | 64.05 | [62.15, 64.46] | 0.39 | 0.960 | 0.46 | 0.079 | 0.25 | 0.263 | 0.21 | [-0.004, 0.008] | 0.71 | 0.786 |
| inferior temporal gyrus | 108.44 | 176.73 | 62.98 | [61.82, 64.13] | 108.44 | 177.38 | 63.57 | [63.04, 65.51] | -1.50 | 0.960 | -0.43 | 0.024 | 0.36 | 0.155 | 0.79 | [0.003, 0.013] | 3.05 | 0.023 |
| precuneus cortex | 153.11 | 251.21 | 64.07 | [62.74, 65.4] | 153.11 | 247.79 | 61.84 | [63.24, 66.17] | -0.63 | 0.960 | -1.77 | 0.000 | -1.37 | 0.000 | 0.40 | [-0.004, 0.012] | 0.96 | 0.786 |
| lateral occipital cortex | 119.09 | 195.19 | 63.90 | [62.6, 65.2] | 119.09 | 193.96 | 62.87 | [62.33, 64.96] | 0.27 | 0.960 | -0.48 | 0.009 | -0.63 | 0.014 | 0.15 | [-0.004, 0.007] | 0.56 | 0.786 |
| lingual gyrus | 125.33 | 207.24 | 65.36 | [63.83, 66.9] | 125.33 | 208.00 | 65.97 | [64.19, 67.15] | -0.28 | 0.960 | 0.18 | 0.440 | 0.37 | 0.268 | 0.19 | [-0.005, 0.009] | 0.53 | 0.786 |
| superior frontal gyrus | 266.60 | 440.84 | 65.35 | [64.13, 66.58] | 266.60 | 446.59 | 67.51 | [63.43, 65.59] | 1.02 | 0.960 | 1.81 | 0.000 | 1.30 | 0.000 | 0.51 | [-0.001, 0.011] | 1.77 | 0.327 |
| middle temporal gyrus | 121.32 | 197.96 | 63.17 | [61.87, 64.47] | 121.32 | 199.32 | 64.29 | [63.02, 65.64] | -1.23 | 0.960 | -0.02 | 0.920 | 0.68 | 0.009 | 0.70 | [0.001, 0.013] | 2.30 | 0.122 |
| paracentral lobule | 60.42 | 99.51 | 64.70 | [63.24, 66.16] | 60.42 | 99.45 | 64.60 | [64.15, 67.13] | -0.88 | 0.960 | -0.63 | 0.015 | -0.06 | 0.877 | 0.58 | [-0.002, 0.014] | 1.38 | 0.565 |
| inferior frontal gyrus | 122.17 | 200.92 | 64.46 | [63.04, 65.87] | 122.17 | 204.46 | 67.36 | [62.64, 65.86] | 0.19 | 0.960 | 1.86 | 0.000 | 1.73 | 0.000 | 0.13 | [-0.004, 0.007] | 0.45 | 0.786 |
| superior parietal cortex | 154.49 | 254.36 | 64.65 | [63.41, 65.89] | 154.49 | 254.63 | 64.83 | [63.37, 65.96] | -0.02 | 0.982 | 0.09 | 0.721 | 0.11 | 0.591 | 0.02 | [-0.005, 0.005] | 0.06 | 0.950 |
| superior temporal gyrus | 165.20 | 271.39 | 64.28 | [62.85, 65.71] | 165.20 | 273.72 | 65.68 | [63.43, 66.01] | -0.46 | 0.960 | 0.59 | 0.024 | 0.86 | 0.013 | 0.27 | [-0.004, 0.01] | 0.74 | 0.786 |
| supramarginal gyrus | 120.39 | 197.91 | 64.39 | [63.16, 65.62] | 120.39 | 196.64 | 63.33 | [61.69, 64.27] | 1.55 | 0.960 | 0.21 | 0.440 | -0.64 | 0.004 | 0.85 | [0.003, 0.014] | 3.00 | 0.023 |
| postcentral gyrus | 168.12 | 276.14 | 64.25 | [63.12, 65.38] | 168.12 | 279.00 | 65.96 | [63.03, 65.14] | 0.21 | 0.960 | 1.14 | 0.000 | 1.04 | 0.000 | 0.10 | [-0.004, 0.006] | 0.40 | 0.786 |

*SP volume**

| | left growth | | | | right growth | | | | left vs right | | AI (%) | | | | | | | |
|---|---|---|---|---|---|---|---|---|---|---|---|---|---|---|---|---|---|---|
| **ROI** | **early** | **mid** | **%growth** | **CI** | **early** | **mid** | **%growth** | **CI** | **Z** | **p (corr.)** | **early AI** | **p (corr.)** | **mid AI** | **p (corr.)** | **ΔAI** | **CI [AI/100]** | **Z** | **p (corr.)** |
| middle frontal gyrus | 6698.81 | 13415.86 | 100.27 | [95.33, 105.22] | 7186.82 | 14228.06 | 97.97 | [95.33, 103.49] | 0.61 | 0.577 | 6.81 | 0.002 | 5.82 | 0.000 | 0.99 | [-0.03, 0.05] | 0.46 | 0.683 |
| cuneus cortex | 3148.51 | 5404.52 | 71.65 | [66.46, 76.85] | 3019.52 | 5105.18 | 69.07 | [66.46, 74.53] | 0.67 | 0.569 | -4.16 | 0.147 | -5.78 | 0.014 | 1.61 | [-0.04, 0.08] | 0.54 | 0.671 |
| precentral gyrus | 6940.22 | 12702.99 | 83.03 | [78.45, 87.62] | 6919.08 | 11438.01 | 65.31 | [78.45, 69.83] | 5.40 | 0.000 | -0.32 | 0.934 | -10.44 | 0.000 | 10.12 | [0.06, 0.14] | 4.91 | 0.000 |
| fusiform gyrus | 2396.05 | 4768.00 | 98.99 | [92.74, 105.25] | 2411.88 | 4911.63 | 103.64 | [92.74, 110.34] | -0.99 | 0.418 | 0.38 | 0.934 | 2.87 | 0.071 | 2.49 | [-0.03, 0.08] | 0.89 | 0.485 |
| inferior parietal cortex | 6717.28 | 12385.97 | 84.39 | [79.85, 88.93] | 6038.96 | 11806.62 | 95.51 | [79.85, 100.44] | -3.25 | 0.004 | -10.87 | 0.000 | -4.80 | 0.000 | 6.07 | [0.01, 0.11] | 2.40 | 0.035 |
| inferior temporal gyrus | 3327.79 | 6882.13 | 106.81 | [101.73, 111.89] | 3382.60 | 6748.54 | 99.51 | [101.73, 104.12] | 2.09 | 0.063 | 1.81 | 0.644 | -1.95 | 0.070 | 3.76 | [-0.01, 0.09] | 1.48 | 0.213 |
| precuneus cortex | 4890.59 | 8263.66 | 68.97 | [64.12, 73.83] | 4198.53 | 6710.08 | 59.82 | [64.12, 64.28] | 2.72 | 0.016 | -15.29 | 0.000 | -20.41 | 0.000 | 5.12 | [0.01, 0.09] | 2.46 | 0.035 |
| lateral occipital cortex | 3954.77 | 7307.88 | 84.79 | [80.36, 89.22] | 3970.11 | 7847.78 | 97.67 | [80.36, 102.68] | -3.77 | 0.001 | 0.27 | 0.934 | 7.00 | 0.000 | 6.73 | [0.02, 0.11] | 3.00 | 0.013 |
| lingual gyrus | 3045.14 | 5663.32 | 85.98 | [81.57, 90.39] | 3175.21 | 6226.24 | 96.09 | [81.57, 100.86] | -3.05 | 0.007 | 4.21 | 0.005 | 9.30 | 0.000 | 5.08 | [0.01, 0.09] | 2.35 | 0.036 |
| superior frontal gyrus | 8057.76 | 15153.82 | 88.06 | [84.56, 91.57] | 7740.81 | 15646.58 | 102.13 | [84.56, 106.52] | -4.91 | 0.000 | -4.18 | 0.043 | 3.13 | 0.003 | 7.31 | [0.04, 0.11] | 3.91 | 0.001 |
| middle temporal gyrus | 4682.51 | 7813.79 | 66.87 | [61.11, 72.64] | 4278.97 | 6955.96 | 62.56 | [61.11, 67.56] | 1.11 | 0.380 | -8.65 | 0.000 | -11.80 | 0.000 | 3.14 | [-0.02, 0.08] | 1.28 | 0.283 |
| paracentral lobule | 1853.78 | 3298.27 | 77.92 | [71.62, 84.23] | 1896.76 | 3669.81 | 93.48 | [71.62, 98.55] | -3.77 | 0.001 | 2.50 | 0.178 | 10.94 | 0.000 | 8.43 | [0.03, 0.14] | 2.94 | 0.013 |
| inferior frontal gyrus | 3423.60 | 6535.41 | 90.89 | [84.99, 96.8] | 3675.06 | 6696.62 | 82.22 | [84.99, 86.8] | 2.27 | 0.043 | 7.26 | 0.011 | 2.71 | 0.099 | 4.55 | [-0.01, 0.1] | 1.66 | 0.165 |
| superior parietal cortex | 5467.88 | 9786.54 | 78.98 | [73.12, 84.84] | 5603.51 | 10599.01 | 89.15 | [73.12, 94.45] | -2.52 | 0.025 | 2.50 | 0.184 | 8.14 | 0.000 | 5.64 | [0.02, 0.09] | 2.89 | 0.013 |
| superior temporal gyrus | 5289.55 | 8421.89 | 59.22 | [52.12, 66.31] | 5254.66 | 8462.13 | 61.04 | [52.12, 66.26] | -0.41 | 0.685 | -0.38 | 0.934 | 1.32 | 0.583 | 1.70 | [-0.04, 0.07] | 0.61 | 0.657 |
| supramarginal gyrus | 4712.92 | 8383.54 | 77.88 | [72.97, 82.8] | 4737.76 | 8283.83 | 74.85 | [72.97, 81.04] | 0.75 | 0.548 | -0.02 | 0.993 | -1.30 | 0.570 | 1.28 | [-0.05, 0.08] | 0.40 | 0.688 |
| postcentral gyrus | 6537.66 | 9146.77 | 39.91 | [34.27, 45.55] | 6683.09 | 9906.64 | 48.23 | [34.27, 54.13] | -2.00 | 0.070 | 2.17 | 0.236 | 7.99 | 0.000 | 5.82 | [0.01, 0.11] | 2.42 | 0.035 |

**Supp. Table 8.** Silhouette scores quantifying the quality of different clustering solutions. Higher scores indicate better separation and cohesion within clusters.

| | *Silhouette score* | | |
|---|---|---|---|
| **Cluster number** | **SP volume** | **SP surface area** | **SP thickness** |
| 2 | 0.655 | 0.709 | 0.465 |
| 3 | 0.361 | 0.406 | 0.507 |
| 4 | 0.357 | 0.423 | 0.513 |
| 5 | 0.331 | 0.408 | 0.575 |
| 6 | 0.356 | 0.438 | 0.603 |
| 7 | 0.357 | 0.430 | 0.559 |
| 8 | 0.350 | 0.419 | 0.556 |
| 9 | 0.358 | 0.368 | 0.514 |
| 10 | 0.360 | 0.385 | 0.463 |
| 11 | 0.343 | 0.367 | 0.451 |
| 12 | 0.328 | 0.347 | 0.460 |
| 13 | 0.327 | 0.352 | 0.451 |
| 14 | 0.319 | 0.354 | 0.467 |
| 15 | 0.288 | 0.342 | 0.506 |
| 16 | 0.280 | 0.354 | 0.481 |
| 17 | 0.263 | 0.335 | 0.453 |
| 18 | 0.260 | 0.367 | 0.457 |
| 19 | 0.241 | 0.322 | 0.443 |
| 20 | 0.229 | 0.314 | 0.459 |
| 21 | 0.209 | 0.291 | 0.446 |
| 22 | 0.194 | 0.298 | 0.453 |
| 23 | 0.190 | 0.284 | 0.461 |
| 24 | 0.182 | 0.266 | 0.410 |
| 25 | 0.169 | 0.262 | 0.380 |
| 26 | 0.147 | 0.246 | 0.365 |
| 27 | 0.132 | 0.216 | 0.316 |
| 28 | 0.118 | 0.202 | 0.280 |
| 29 | 0.107 | 0.172 | 0.236 |
| 30 | 0.085 | 0.130 | 0.197 |
| 31 | 0.067 | 0.107 | 0.157 |
| 32 | 0.048 | 0.098 | 0.113 |
| 33 | 0.048 | 0.098 | 0.113 |

optimal number (highest Silhouette score)

final cluster number
